# De-Novo Designed Antibacterial N95 Facial Mask: Comprising a Nano-Garden Using ZnO Nanoflower

**DOI:** 10.64898/2026.04.20.719592

**Authors:** Priyanka Bhadra, Rebanta Roy, Subhrangsu Chaterjee

## Abstract

Nowadays N95 facial mask has gain huge attention due to COVID19 pandemic situation and it serves as the prime PPE. Though the microbes can be restricted to get inside the human body due to the presence of mask temporarily, but over the time, bacteria and other microbes may get entrapped into the threads of the mask itself and thus acting as a storage chamber of microbes. It is necessary to eliminate them from the mask surface. To do so different floral structured Nano-ZnO with variable oriented arrangement of petals were fabricated on the surface of the N95 mask and further characterized through instrumentations including XRD, FTIR,UV-Vis, Fluorescence-Spectroscopy, SEM, DLS. The average crystallite size calculated for synthesized four different ZnO nanoflower were 25.19 nm, 23.46 nm, 27.27 nm and 31.78 nm (for **glycerol**, **PEG**, **EDTA**, **Chitosan** assisted) respectively. The antimicrobial activity was investigated by standard microbial broth dilution assay and Kirby-Bauer test which assured the inhibition of the bacterial growth. The MIC-MBC value of ZnO nanoflowers for *E.coli* and *B. subtilis* were found to be effective at dilution of 250 µg/ml and 100 µg/ml. Additionally a modified Kirby-Bauer assay has been designed to investigate the killing efficiency of the bacteria (*E.coli*).

Fig. - Graphical Abstract

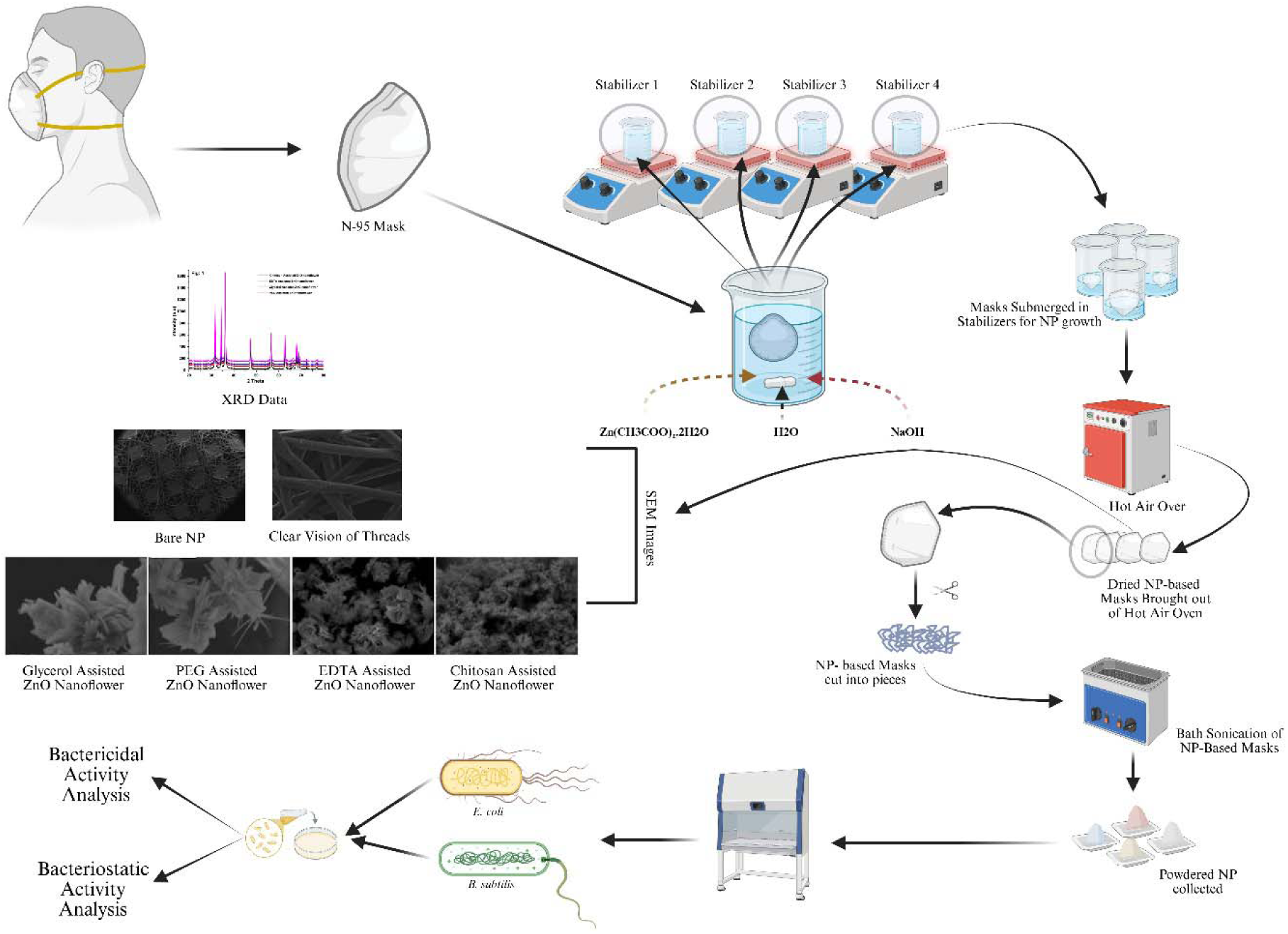

## Introduction

As humans are marching forward with the civilization, diseases are becoming deadlier with unique levels of adaptations of the pathogenic micro-organisms. In the recent years scientists have valued the pathogenicity of these microorganisms and have hence tried to bring up novel therapeutic strategies. In the relatedness to the microbes, personal protective equipments (PPEs) play important roles in safeguarding and protecting the human body from potential contamination from microbes [1]. COVID19 is one of the recent examples of encounters of humans with the microbial world [2]. The whole world witnessed how the PPEs saved the ones who were first line warriors against this fatal viral disease. PPEs are available in different forms, ranging from whole body protective equipment to an equipment covering a local body part [3]. One of the best and easily available form of PPEs are face masks. The face masks are of many types ranging from low grade protection to high grade protection viz., scarves and fabric masks, surgical masks, hygienic masks, KN95 masks, N95 masks, FFP1 masks, FFP2 masks, FFP3 masks, etc. The most widely available protective form of masks are N95 (USA Standard) face masks which gives around 95% protection to particles which are of the size 0.3 microns or more [4], including bacteria and viruses especially the SARS CoV2 in the recent context. The N in N95 name comes from “non oil”, which suggests that N95 masks can be used only in the presence of non oil particulate matters [5]. The N95 masks are made of non woven polypropylene layers. These closely arranged polypropylene layers increase the compactness in the masks making the masks much more resistant to particulate matters [6].

Now, there have been various reports which states that N95 masks lose their resistance to microbes after few rounds of washing, due to the loss of compactness in the polypropylene layers [7]. So, the modification of such a property becomes so important in terms of both durability and cost efficiency of an N95 mask. Also with subsequent use, there might be a chance of growth of microbial films on both the inner and outer surfaces of the mask [8]. In the recent years several forms of research proved that the adverse health effect and mortality related to COVID19 was not only due to the symptoms arising from the infection of the SARS-CoV2 virus itself, but the majorly due to the secondary forms of infections arising alongside COVID19 [9]. The secondary or tertiary infections were mostly associated with bacterial organisms. Hence, the term “COVID19 associated disorders” became so popular [10]. Even the long term effect on health due to COVID19 has been reported due to the massive health impact caused by secondary and tertiary infections [11]. Reports also state that COVID19 infection, helped other disease associated microbes, which were already present in our body, kept in an inactive or dormant state by the immune system, to grow by deceiving the immune response [12]. This can be suspected to be one of the major causes behind healthy people falling severely ill due to COVID19. This was common for healthy people, non patients, patients staying under home isolation and for patients staying in hospitals. The use of same masks repeatedly with continuous washing at frequent intervals could be one of the reasons for the increased prevalence of the secondary infections. Hence, the masks may lose the property of microbial resistance slowly and the chances of secondary and tertiary microbial growth on it’s surface also increases by many folds. So, using N95 masks as a surface for the growth of potential antimicrobial nanoparticles may be a potent solution to this.

The recent encounter of humans with the masks were during the COVID19 pandemic, when WHO, as well as the Governments of various countries and federal states across the whole world made masks compulsory in public places. After that period masks has been still a daily life commodity of many people around the globe in their day to day life. Even in case of laboratory, both research and industry as well as in case of hospitals, masks are a commodity for daily use. So, the importance of providing microbe free masks become very important.

In this regards nanomedicine, which is a branch of nanosciences has given new signs of hope in disease diagnosis and improvement in treatment regime. Many nanoparticles and nanomaterials have the property to resist microbial growth, Zinc Oxide (ZnO) Nanoparticles are such a type of nanoparticle which has shown exclusive resistance to microbes [13][14]. Such, microbe sensing resistance of these NPs has brought newer dimensions into antimicrobial research. ZnO has been proven to show a very low toxicity, which makes it a potential candidate to be used as a bio based nanomaterial [15]. Also the characterization of the ZnO nanoparticles post synthesis is not so tedious. The basic biophysical methods, for example UV Spectroscopy, XRD, SEM, FT-IR, DLS and fluorescence spectroscopy can be highly instrumental in characterizing the nanoparticles [16]. In many cases ZnO NPs are used as carriers of different types of drugs inside the human body due to the unique meshwork and surface chemistry which the particular nanomaterial forms [17]. Again due to the surface charge, these ZnO nanoparticles have also been used as potential alternative to semiconductor materials, solar cells as well as photocatalysts[18]. But what has surprised most researchers is their antimicrobial effect which it shows, where ZnO NPs have been found to have profound antimicrobial impact on several biofilm forming heterogenous pathogens [19][20]. In our work we have tried to utilize this unique microbe sensing resisting property of the ZnO Nanomaterial to develop a novel therapeutic approach to tackle microbial growth on different surfaces which play major roles in daily life usage. The grafting of nanostructures onto polypropylene based N95 mask surface is gaining a crucial part of research which helps to transform the mask from a simple physical filter into a biocidal protective shield. As the polypropylene is chemically inert, the grafting process requires specific techniques to achieve durable attachment of the differential nanostructures over the mask surface [21][22]. In our work, we have used various capping agents viz., Chitosan, EDTA, Glycerol, PEG to the base nanomaterial to improve the process of nucleation and oriented growth towards a specific direction, since capping has been found to be instrumental in causing improved rates of reaction between the Nanomaterial and their media of synthesis. In our work we have looked to counter both the routes of limiting bacterial population increment, firstly the bacteriostatic activity, where we have looked to counter the bacteria by limiting the growth using the different morphological ZnO Nanomaterial, on the other hand we have also looked for killing the bacteria using those ZnO Nanomaterial by using the bacteriocidal activity. Though our work over here is limited in checking bacteriostatic and bacteriocidal activity, but in future we are looking forward to test the antifungal and antiviral properties of the Nanoflower based masks. In our novel work we have used a single step process instead of a multistep process based on differential ZnO nanostructures. Also, the re-usability of the N95 masks increased further even after washing them several times with both hot and cold water, increasing the novelty further to some extent.

## Method

### Materials

Zinc Acetate dehydrate 99.5% extra pure from SRL, Chitosan powder (Medium molecular weight) 50-500m, Pas, 90% DA (Deacetylated Chitin extrapure) were purchased from Millipore Sigma and used as received. Polyethelene Glycol 200 (PEG) was purchased from Merck, Acetic acid glacial (GAA, 99.5% extra pure) was purchased from LOBA CHEMIE PVT LTD, Zinc sulphate monohydrate (99.9% trace metals basis), Amonium Hydroxide solution (25% NH_3_ basis) were purchased from sigma, EDTA Dipotassium Salt (Ethylenediaminetetraacetic Acid Dipotassium Salt) pure,98%, Glycerol (99.5% from sigma) and Sodium Hydroxide Pellets (from MERCK) were purchased. Commonly used N95 respiratory mask was purchased from local medical shop. *Escherichia coli* (ATCC 25922) was gifted as a lyophilized powder from the Department of Microbiology, Bose Institute Kolkata. *Bacillus subtilis* (Ehrenberg) Cohn (ATCC - 6051) was gifted by Department of Microbiology, Bose Institute Kolkata as a lyophilized powder. Luria Bertini (LB) broth, agar, and phosphate buffered saline (PBS) were purchased from Millipore-Sigma.

There were two kinds of process methods have been followed in this current research work i.e In-situ synthesis of different structured ZnO nanomaterial over the N95 respirator mask and Antimicrobial activity study including both Bactericidal and Bacteriostatic activity. There were four different kinds of ZnO nanomaterials have been grafted over the N95 respirator mask. The process of functionalizing the polypropylene (PP) fibers of an N95 mask with several decorating nanomaterials is typically achieved through a multi-step coating or grafting technique, with the main goal of adding powerful antimicrobial properties without compromising filtration. Since polypropylene is chemically inert, direct covalent grafting mechanism is not possible until the PP surface is not activated first by treated using methods like Plasma Activation (e.g., using O_2_ or Ar plasma) or chemical oxidation. This introduces reactive functional groups (like hydroxyl or carboxyl groups) onto the non-polar PP surface. In the current project we apply Glycerol, PEG, EDTA and Chitosan based ZnO nanostructure functionalisation to the non-woven polypropylene fabric of a mask by simple adhesion where binding occurs primarily through physical adsorption and interfacial forces (like van der walls forces) between the nanostructure and the PP surface. It involves a multi-step process, typically combining the synthesis of the nanomaterial with its application onto the fiber substrate. In the standardized procedure, combining all these nanocomponents onto the mask thread requires established techniques for nanoparticle synthesis (often a wet chemical method like precipitation or sol-gel) and fiber functionalization. In this current work we have chosen one pot synthesis instead of multistep process where the nucleation and growth of the differential ZnO nanostructures initiates directly from the precursor salts along with the individual stabilizing agent and the entire fabrication take place in the submerged condition by using polypropylene thread as a template. Precursor, zinc acetate dihydrate, Zn(CH_3_COO)_2_. 2H_2_O dissolution in deionised water can produce Zn^2+^ ions in presence of various stabilizing agents. EDTA acts as a chelating agent, controlling the zinc ion concentration and reaction kinetics, which is crucial for uniform size distribution and inhibiting premature crystal growth. Whereas PEG acts as a capping/stabilizing agent and a template, preventing agglomeration of the growing ZnO nanoparticles *via* steric hindrance and controlling the size/shape. Polymer Chitosan and Glycerol can act as another stabilizing agent. Incorporation of NaOH, a reducing agent causes the Zn^2+^ ions to react and form zinc hydroxide Zn(OH)_2_ or zinc precursors. Heating the solution drives the dehydration and crystallization process, leading to the formation of the Wurtzite crystal structure of ZnO nanoparticles. The coated thread/fabric is dried, often with mild heating, to remove the solvent and firmly anchor the stabilizing agent/ZnO nanoflower layer onto the non-polar polypropylene surface. In a brief, 0.5 M Zinc acetate solution was dissolved in 500 ml of deionised water (DI). 2% of individual solution of Glycerol, PEG, EDTA and Chitosan with Glacial acetic acid were inserted into the Zinc acetate solution separately. The solution mixture was agitated for 2 hrs in warm condition (40° C). Individual N95 respirator mask was dipped in four different solution mixture and put it in the Orbital shaker at 200 rpm for 2 hours. This solution treated masks were dipped again into 1M NaOH solution for another 30 mins in the orbital shaker at the same rpm rate followed by Deionised water wash for three times. Finally individual treated mask was kept in an oven and dried at 50 °C until it become dry completely.

### Nanomaterial phase recovery from N95 respirator mask

Different structured ZnO nanomaterials were recovered from the mask by shaking in an orbital shaker for 15 mins at 200 rpm in 10 mL of sterile phosphate buffered saline (PBS) followed by ultra-sonication at about 15 mins. As our treated mask was prepared by completely submerging into the precursor solution from where the nanomaterial can be originated and deposited over the polymer threads of the mask, the nanomaterial need to be extracted out from the layer. Shaking performed a higher recovery efficiency than vortex, so shaking was used in most of the experiments in this study. It was highly recommendable to carry out further recovery of the nanomaterial from the layer by ultra-sonication [23].

### *E.coli* and *B.subtilis* strain, medium and cultivation

The revived ampoule of *Escherichia coli* (ATCC 25922) bacteria was centrifuged at 11,000 rpm for 5 min at 4 degree C in LB broth. 20 ml of sterile normal saline was added after discarding the supernatant. The remaining viable bacterial precipitate was maintained as stock. Then, the concentration was adjusted by spectrophotometer to an optical density of 0.10 at 600 nm (10^8^ CFU/ml, 0.5McFarland’s standard). Simultaneously the Bacillus bacterial strain was also cultured in the same way as like as the *E.coli* strain.

### Bacteriostatic activity study (Minimum inhibition concentration determination)

The antibacterial efficacy of different structured ZnO nanomaterial was observed by the standard broth dilution or microdilution method (CLSI M07-A8) where the visible growth of microorganisms in the agar broth have been noticed. ZnO nanomaterial in different concentrations i.e. 1mg/ml, 750 µg/ml, 500 µg/ml, 250 µg/ml, 100 µg/ml and 1 µg/ml with adjusted bacterial concentration (10^8^ CFU/ml, 0.5 McFarland’s standard) were used to determine MIC in 20 ml of LB broth. Simultaneously the marketed antibiotic drug named as Amoxicilin and erythromiocin was used as positive control, inoculated broth was treated as control and simple Deionised water was marked as negative control. These all culture tubes were incubated for 24 hr at 37 ° C. The MIC value is the lowest concentration of ZnO nanomaterials where no visible growth is seen in the tubes. The visual turbidity of the tubes were noted, both before and after incubation to confirm the MIC value. Additionally the antibacterial study of different structured ZnO nanmaterial was studied using the well diffusion/agar diffusion or cup plate (Kirby Bauer) method against gram negative *E.coli* bacteria as well as gram positive *B.subtilis* bacteria where the 100 µl bacterial solution was plated on the 20 ml LB agar plate. After solidifying the agar plate, small cup was scooped out from the each plate. Here also the standard antibiotic Amoxycilin, erythromyocin and pure Deionised water were chosen as the control group and various concentration of ZnO nanmaterials were treated as unknown samples. 100 µl of individual test sample solution from each culture tube was withdrawn and put it into the scooped cup created on the each agar plate to measure the zone of inhibition. The bacteriostatic activity of individual test sample was parallely examined by using 0.5 mg/ 20 ml broth in an orbital shaker at 200 rpm and incubated in 37 ° C in dark condition. In each culture tube, 100 µl of *E.coli* bacterial solution (OD 0.10 at 600 nm) was kept and checked the bacterial cell doubling by evaluating the OD measurement at 600 nm wavelength with time interval ranges from T0, T1……..T24 (T0-Time 0 hr). The bacterial growth was measured using ND-1000 v 3.3.1 spectrophotometer, (Nanodrop Technologies, Inc., Wilmington, USA). Control (no addition of nanomaterial) was also maintained. The total viable count, indicating the number of bacteria that survived after interaction with the different structured ZnO nanomaterial was enumerated by plate count method.

### Minimum bacteriocidal concentration (MBC determination) study

After the MIC determination of the different structured ZnO nanomaterials, aliquots of 50 µl from all the tubes which showed no visible bacterial growth were seeded on LB agar plates and incubated for 24 h at 37 ° C. When approximately all of the bacterial population is killed at the lowest concentration of an antimicrobial agent, it is termed as MBC endpoint. This was done by observing pre and post-incubated agar plates for the presence or absence of both *E.coli* and *B.subtilis* bacteria. A modified version of both bacteriostatic and bacteriocidal activity study was performed using broth culture technique. 100 µl of bacterial solution was plated in the 20ml of LB agar plate and incubated at 37 ° C in dark condition to form the bacterial colony over the agar plate. After 24 hrs a specific portion of cup was scooped out from the bacterial culture plate and 100 µl of different structured nanomaterial solution was poured inside the hole. Nanomaterial loaded bacterial culture plates were further incubated at 37 ° C in an incubator. Upon incubation the nanomaterial solution was diffused inside the LB agar gel to a certain portion and a specific zone of inhibition was formed around the hole. One loop full of bacterial culture was withdrawn from both the inside and outside of the inhibition zone and reloaded into the freshly prepared LB broth (20 ml) to check the bacterial growth in terms of OD (optical density) measurement at 600 nm wavelength with time interval. Both UV-Vis spectrophotometer and Nanodrop were used to optimise the bacterial growth.

### Characterisations of ZnO nanomaterial

XRD analysis of different structured ZnO Nanomaterial: The synthesized differential ZnO nanomaterials were characterised by X Ray Powder Diffraction (XRD, Rigaku, Ultima III, Cu Kα radiation) method in the range of 2θ values within 10°-80° at a scan rate of 5°/min. Both Nanomaterial grafted N95 mask and recovered Nanomaterial from the mask were used for XRD characterising to confirm the Zincite phase purity. The d values were computed and compared with the standard d values of ZnO (JCPDS no: 036-1451) resulting in excellent matching.

### UV-Vis Spectroscopy analysis

The optical transmittance spectra was taken in the UV-Vis range by using UV-Vis Spectroscope (Cary60 Agilent). The band gap was calculated from the absorbance at the wavelength where the four individual ZnO nanomaterial was absorbed most. The calculated band gap values of all nanomaterials were compared with respect of the band gap of the bulk ZnO (□3.23 eV).

### FTIR Spectroscopy analysis

The FTIR spectra were recorded on a Schimadzu FTIR spectroscope (IRPrestige21, 200 VCE) in absorption mode with 50 scans at a resolution of 4 cm^-1^. The all recovered differential ZnO nanomaterials from the N95 respirator mask were mixed with potassium bromide (KBr) and the fixed scan range was in between 400-4500 cm^-1^. FTIR spectroscopy analysis is the process where we can investigate the formation of ZnO nanomaterial with a definite bond formation and at the same time we observe the bonding of the surface modifier molecules to the ZnO surface.

### Morphological study by SEM analysis

Scanning Electron Microscopy is one of the most valuable characterising tool to study the Nanomaterial size and structure. The signal derived from the interaction in between the sample and electron reveals the information about the sample morphology. The size and morphology of the as prepared ZnO samples were carried out by Quanta 200, FEI.

### Dynamic light scattering (DLS) studies

Dynamic light scattering (Malvern Zeta NanoS ZEN2000) is known as photon correlation spectroscopy or quasi-elastic light scattering. This is emerged as simple table-top technique executable under ordinary lab environments to investigate the hydrodynamic size. When DLS is determined to study the nanoparticle size, an important point has to keep in mind which is particle concentration. When the particle concentration is too low, the scattering from the particles is weak and the measurement results are noisy. When the particle concentration is too high, the measurement results are distorted due to particle-particle interactions. These interactions can be particle-particle collisions at high particle concentrations. Or, the interactions can be long range electrostatic interactions. In either case, these interactions change particle motion compared to the motion assumed when calculating particle size from DLS data. At high particle concentration, the data seems normal, but the size results are distorted. Therefore, when developing methods, it is important to keep particle concentration in mind.

## Result and Discussions

Fig 1 shows the diffraction peaks of ZnO Nanomaterial at 100, 002, 101, 102, 110, 103, 112 which correspond respectively to the values in degrees (2θ) at 31.34°, 34.50°, 36.32°, 47.60°, 56.68°, 62.94°. High diffraction peaks indicated the crystalline nature of the material. The crystallite size of the ZnO nanostructure was calculated by Debye Scherrer equation. The high intensity peak at (101) was used to determine the lattice parameters.

**Fig 1:**
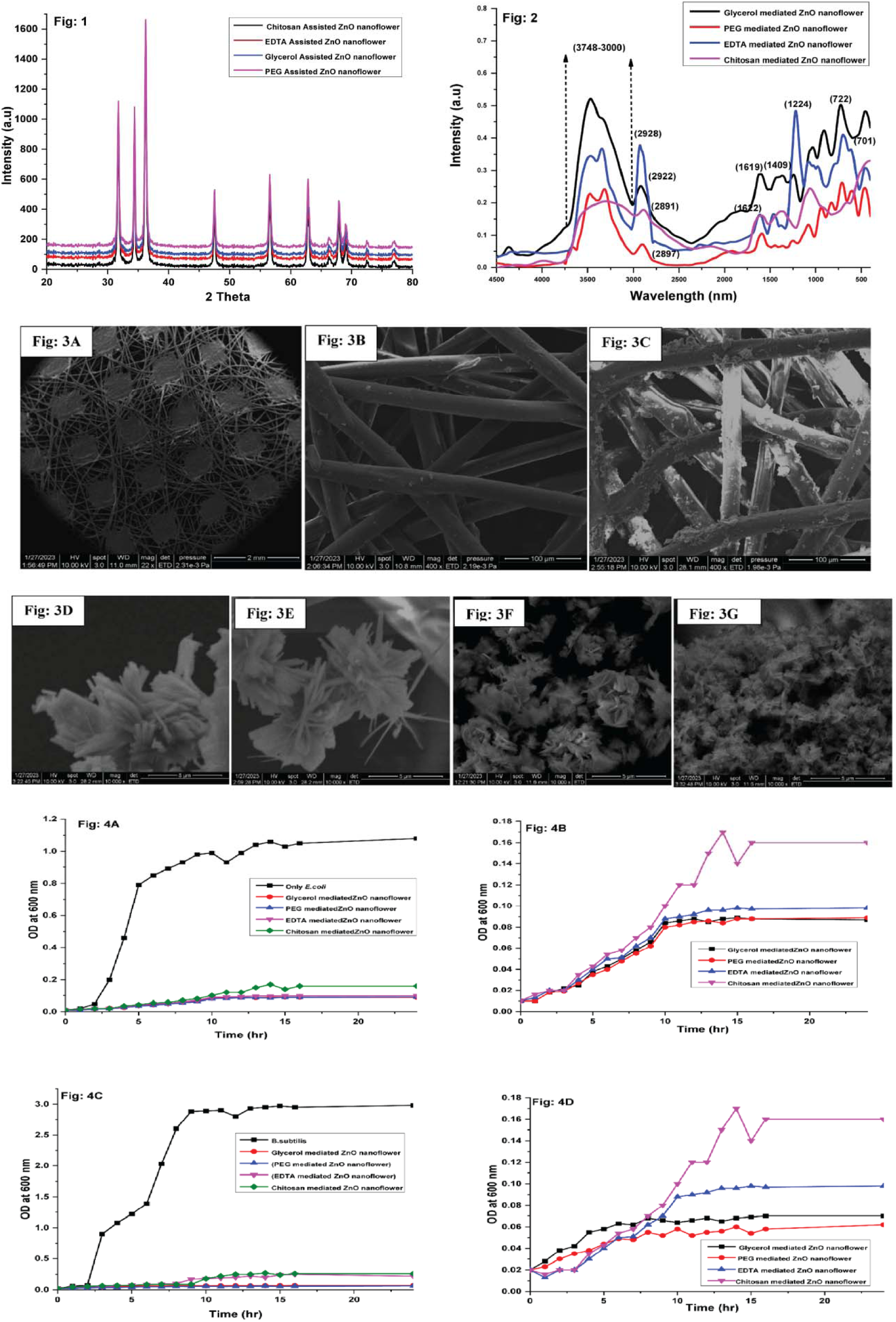
XRD analysis of differential surface modified ZnO nanoflower Fig 2: FTIR absorption spectra of four different patterned ZnO nanoflower Fig 3: SEM image analysis reveals the formation of different patterned ZnO nanoflower over the thread of the N95 mask. **3A** denotes the bare N95 mask without any Nanomaterial deposition; **3B** denotes the clear vision of individual threads without any Nanomaterial deposition (Zoom version); **3C** shows the presence of Nanomaterial attachment over the individual thread; **3D** shows the presence of Nanomaterial attachment over the individual thread (Zoom version); **3E** denotes the high-resolution image of Glycerol assisted ZnO Nanoflower over the N95 mask thread**; 3F** denotes the high-resolution image of PEG assisted ZnO Nanoflower over the N95 mask thread; **3G** denotes the high-resolution image of EDTA assisted ZnO Nanoflower over the N95 mask thread and **3H** denotes the high-resolution image of Chitosan assisted ZnO Nanoflower over the N95 mask thread. **Fig 4A**: Growth curve of *E.coli* bacteria from initial hour to 24 hrs with one hr time interval with and without treatment of four different ZnO nanoflower, **4B**:Growth curve of E.coli bacteria in presence of four different ZnO nanoflower, C: Growth curve of *B.subtilis* bacteria from initial hour to 24 hrs with one hr time interval with and without treatment of four different ZnO nanoflower, D:Growth curve of *B.subtilis* bacteria in presence of four different ZnO nanoflower.

D=[0.9λ/βCosθ]*100

Where D is the size of the ZnO crystallite; λ is the wavelength of Cu Kα radiation at 1.5418 Å; θ is the Bragg diffraction angle, and β is the full width at half maximum intensity of the diffraction peak of the sample. The crystallite size can be measured more accurately by high resolution X-ray diffraction (HRXRD) using the Bond method, which increases peak resolution to find the values of the Lattice parameters. In this study, we determined by XRD powder that most synthesized ZnO crystallites sizes are variable. Lattice constant is the term which is dependent on crystallite size.

hkl2dsinθ=nλ=d_hkl_=λ/2dsinθ

1/d_hkl_^2^=h^2^/a^2^+k^2^/b^2^+l^2^/c^2^

d_hkl_ = a/√h^2^+k^2^+l^2^

a = d_hkl_√h^2^+k^2^+l^2^

n is the diffraction order, for 1^st^ order n=1

λ is the X ray wavelength, Cu Kα radiation at 1.5418 Å

d is the interplanar spacing in Å

### θ is the Bragg diffraction angle

The calculated d values for four different nanomaterials were computed and compared with the standard d values of ZnO (JCPDS NO: 036-1451) and it has been shown the excellent matching in all samples with the extreme purity of being the single crystalline Zincite phase without any other impure phases. The peak broadening in the XRD peaks clearly indicates the formation of nanocrystals. Both the computed and standard d values of ZnO along with the other parameters have been summarised in Table 1. The average crystallite size calculated for synthesized four different ZnO nanomaterial was 25.19 nm (sample 1-S2, Glycerol assisted ZnO nanomaterial), 23.46 nm (sample 2-S3, PEG assisted ZnO nanomaterial), 27.27 nm (sample 3-S5, EDTA mediated ZnO nanomaterial) and 31.78 nm (sample 4-S6, Chitosan capped ZnO nanomaterial). The crystallite size is not necessarily the same as particle size because it assumed to be the size of a coherently diffracting domain. Here it has been use the term “crystallite size” instead of “particle size” [24, 25].

**Table 1:** Computed and standard d values of ZnO along with the other parameters.

| Sample 1-S2, Glycerol mediated ZnO nanoflower |  |  |  |  |  |  |
| --- | --- | --- | --- | --- | --- | --- |
| {hkl} | Peak position | $d_{\text{XRD}}$ (Å<sup>°</sup>) | $D_{\text{JCPDS}}$ (Å<sup>°</sup>) | FWHM (Degree) | Crystallite size D (nm) | Average size D (nm) |
| 100 | 31.34 | 2.8200 | 2.816 | 0.3300 | 25.0028775 | 25.19 |
| 002 | 34.50 | 2.6053 | 2.602 | 0.2438 | 34.1199898 |  |
| 101 | 36.32 | 2.4806 | 2.476 | 0.3360 | 24.88315022 |  |
| 102 | 47.60 | 1.9128 | 1.911 | 0.3291 | 26.38306228 |  |
| 110 | 56.68 | 1.6274 | 1.626 | 0.3989 | 22.62748049 |  |
| 103 | 62.94 | 1.4790 | 1.477 | 0.3976 | 23.42629492 |  |
| 112 | 67.89 | 1.4012 | 1.407 | 0.4806 | 19.92581752 |  |
| Sample 2-S3, PEG mediated ZnO nanoflower |  |  |  |  |  |  |
| 100 | 31.75 | 2.8810 | 2.816 | 0.3533 | 23.37756065 |  |
| 002 | 34.4 | 2.6432 | 2.602 | 0.2667 | 31.18186443 | 23.46 |
| 101 | 36.25 | 2.4931 | 2.476 | 0.3812 | 21.92829258 |  |
| 102 | 47.55 | 1.935 | 1.911 | 0.3682 | 23.57684955 |  |
| 110 | 56.55 | 1.6355 | 1.626 | 0.4012 | 22.48401885 |  |
| 103 | 62.85 | 1.4921 | 1.477 | 0.4112 | 22.64061831 |  |
| 112 | 67.9 | 1.4544 | 1.407 | 0.5016 | 19.09272425 |  |
| Sample 3-S5, EDTA mediated ZnO nanoflower |  |  |  |  |  |  |
| 100 | 31.84 | 2.8101 | 2.816 | 0.3100 | 26.64884 | 27.27 |
| 002 | 34.23 | 2.612 | 2.602 | 0.2341 | 35.50788 |  |
| 101 | 36.55 | 2.4623 | 2.476 | 0.3012 | 27.77644 |  |
| 102 | 47.85 | 1.902 | 1.911 | 0.3003 | 28.94122 |  |
| 110 | 56.68 | 1.622 | 1.626 | 0.3512 | 25.70075 |  |
| 103 | 62.88 | 1.442 | 1.477 | 0.3668 | 25.38526 |  |
| 112 | 67.76 | 1.415 | 1.407 | 0.4563 | 20.97096 |  |
| Sample 4-S6, Chitosan mediated ZnO nanoflower |  |  |  |  |  |  |
| 100 | 31.84 | 2.815 | 2.816 | 0.2514 | 32.86054 | 31.78 |
| 002 | 34.23 | 2.612 | 2.602 | 0.2012 | 41.31409 |  |
| 101 | 36.55 | 2.477 | 2.476 | 0.2611 | 32.04237 |  |
| 102 | 47.85 | 1.926 | 1.911 | 0.2635 | 32.98311 |  |
| 110 | 56.68 | 1.632 | 1.626 | 0.3119 | 28.93909 |  |
| 103 | 62.88 | 1.443 | 1.477 | 0.3012 | 30.91405 |  |
| 112 | 67.76 | 1.412 | 1.407 | 0.4077 | 23.4708 |  |

The UV-VIS absorption spectra of four different ZnO nanostructures were recorded as shown in **Supplementary 1 (S1)** file. According to this image, the absorption maxima of the individual sample became changed in terms of plot intensity, plot shifting and plot nature. In sample 1, glycerol mediated ZnO nanomaterial shows continuous broad spectrum of absorption with a maxima at 356 nm. Sample 2 (PEG assisted) and sample 3 (EDTA assisted) shows the absorption maxima at 340 nm and 363 nm. Whereas the chitosan assisted sample 4 indicates a different kind of absorption spectra where the absorption maxima shows at two variable positions such as 360 nm and 420 nm respectively. Compared with bulked ZnO material, which shows absorption maxima around 375 nm, all four of the ZnO nanomaterial samples have shown obvious blue shift due to quantum confinement. The band gap arises due to the transition of electrons from valence band to conduction band, indicating the shifting of the band gap. The integration of ZnO nanomaterial leads to an enhanced band gap due to the formation of equilibrium in the ZnO nanomaterial Fermi level. The bulk ZnO material band gap is 3.23 eV. The formula was used to compute the bandgap energy (Eg) of ZnO nanoparticles.

E_g_ = hc/λ

The calculation of the corresponding band gap by using Planck’s constant (h), the velocity of light (c), and the wavelength of UV (380×10^-9^ m) resulted in an observed band gap of 3.23 eV. **Supplementary 2 (S2)** shows the optical band gap graph which is plotted by using a Tauc plot technique of four individual ZnO nanomaterial. The absorbance was taken along the vertical axis and the energy was taken along a horizontal axis which was in electron Volt (eV). The straight line was drawn which was maximum touching the graph and pointing at a point which was corresponding as band gap of the material. The band gap energy estimated from the absorption spectrum of four different ZnO nanomaterial was ∼3.45 eV, ∼3.71 eV, ∼3.49 eV and (∼3.42 eV along with ∼ 2.89 eV) which was higher than the standard value of ZnO (3.23 eV). Based on the band gap estimation values, it indicated the fact that upon enhancement of the ZnO nanomaterial band gap with respect of bulk material band gap, the sizes of the material tend towards nano range. It was observed that the energy band gap is size dependent and increases with a decrease in crystallite size of ZnO nanostructure due to the optical confinement effect. It was also revealed that the band gap of the material was not only affected by the size of the material itself, morphology as well as the synthesis methodologies have the impactful effect on it. The behaviour of the band gap could be explained on the basis of surface defects and surface adsorbed species. Here in the synthesis process, there were four different individual surface adsorbed/assisted molecules have been used which might be the reason of variable band gap values with respect of bulk material [26,27].

**Fig 2** depicted the FTIR absorption spectra of the glycerol assisted ZnO nanomaterial revealed that the broad peak appeared in the range between 3748–3000 cm^-1^ wavenumber. This was due to the presence of hydroxyl (O–H) groups. Spectra at 2922 cm^-1^ corresponds to CH_2_ strechning, which came from glycerol backbone. The hydrogen bond splitting was observable at 1619 cm^-1^ and 1409 cm^-1^ respectively [28]. The peak at around 722 cm^-1^ is attributable to Zn-O stretching vibration mode. In PEG assisted ZnO nanomaterial, there were three individual peak originated at 3672 cm^-1^, 3469 cm^-1^ and 3316 cm^-1^ in the range between 3748–3000 cm^-1^ wavenumber which corresponded the hydroxyl (O–H) groups. The sharp absorption peak at 2897 cm^−1^ was due to the C−H stretching band and the peak noticed at 1359 cm^−1^ and 1448 cm^−1^ corresponds to the C−O−H bending vibration. The peak at 1079 cm^−1^ was attributed to the C−O stretching vibration. The observed peak at 831 cm^−1^ and 939 cm^−1^ corresponds to the long polymeric chain bending in the polymer PEG. The Zn−O stretching mode at 710 cm^−1^ and 596 cm^−1^ in the ZnO spectrum was present in the PEG assisted ZnO nanomaterial [29]. The bonding nature of EDTA with ZnO nanomaterial was investigated by FTIR spectra where it depicted the intense band at 3475 cm^−1^ and 3348 cm^−1^ is attributable to O-H stretching vibration mode which is probably hydrogen-bonded O-H stretching mode in chelates i.e. Zn^2+^-EDTA complex. The free O-H stretching vibration occured above 3600 cm^-1^ which was already observed in PEG assisted ZnO sample. Spectra at 2928 cm^-1^ corresponds to CH_2_ strechning, which came from EDTA backbone [30]. The sharp strong absorption peak at 1622 cm^-1^, confirming the bonding of EDTA on the surface of ZnO by chemisorption of carboxylate group. The absorption peak at 1224 cm^-1^ was attributable to -C-N stretching mode, which confirms the capping of EDTA on the ZnO surface. The Zn−O stretching mode observed at 701 cm^−1^. Chitosan assisted ZnO nanomaterial also confirmed the bonding of nano-ZnO with the chitosan polymer chain which is already well matched with our previous paper. [31] The peak at 3319 cm^-1^ and 2891 cm^-1^ corresponded the hydroxyl (O–H) groups and CH_2_ strechning, which came from Chitosan, a polysaccharide backbone. The band at 1614 cm^-1^ corresponded to the Amide I (C=O) whereas the band at 1375 cm^-1^ corresponded to the Amide III (C-N) bond in chitosan assisted sample. The Zn−O stretching mode observed at 665 cm^−1^ which has confirmed the bonding of ZnO with the Chitosan backbone.

The effects of the different molecules such as PEG, glycerol, EDTA and chitosan in the precursor solution of the nanomaterial samples on the morphology and variable shape of the ZnO nanomaterials were shown in **Fig 3**. The figure showed the powder SEM images of the resultant products where it revealed that the synthesized nanostructures have a differential flower like morphology. SEM micrographs were clearly indicated that the nanoflowers were three dimensional in appearance with several nanopetals at about 40 nm in width and several microns in length. The SEM micrograph of nanoflower was clearly portrayed that the nanopetals were actually the rod like morphology which arises from the centre of the nanoflower. To better understand the basic structure of the emerged nanoflowers which were budded from the baseline of the N95 mask thread, the synthesised nanoflowers were sonicated in bath sonicator for 10 mins and subsequently micrograph was taken which shows the formation of nano-flower composed of several nanorod like structures. These nanoflowers were actually originated from the rod like structure which means flowers are the resultant of agglomeration of rods in a systematic manner to ultimately formed nanoflower morphology. An interesting thing was seen in all those synthesized nanoflowers where all the sample’s morphology was unique as flower like structure with variation in petals arrangement which finally leaded to form differential floral pattern. **Fig 3A** depicted the SEM image of three layers of N95 mask which was composed of three different individual layer. Both the inner and outer layer made of spunbond PP fibers and middle layer made of melblown fibers. In a control N95 mask **(Fig 3B),** the all threads of the outer layer seems to be transparent and clear without showing any existence of nanomaterial deposition whereas the nanomaterial treated mask **(Fig 3C-3H)** shows bounded molecules throughout the entire surface of the mask outer layer. It has been postulated that upon deposition of precursor solution of ZnO nanomaterial in presence of several growth enhancer molecules, there had been formed an initial ZnO seed layer on the surface of the N95 mask. ZnO nanorods can grow radially and perpendicularly to the seed layer, resulting in the formation of ZnO nanoflowers. As the seed layer was patterned in an array structure, the resulting ZnO nanoflowers also exhibited an array pattern. When the *insitu* ZnO nanomaterial was synthesised by glycerol assisted medium, the rod like structure became thicker and all of the rod shapes were emerged from a single baseline i.e. ZnO seed layer **(Fig 3E).** In case of PEG assisted ZnO nanomaterial sample, the crystalline ZnO nanorods were oriented in different pattern which leaded to a new floral formation. Here in this case the flower petal which was actually the nanorod became sharp and elongated with the size range of 2-3 µm in length and 40 nm in thickness **(Fig 3F).** EDTA capped ZnO nanomaterial was very unique in shape. The floral petals were arranged here in such a way to form a flower which looks alike marigold. The flowers were very compact in structure and uniformly distributed over the entire mask. There was a gap in between two adjacent flower **(Fig 3G).** Whereas this floral pattern became same with a little variation where the flowers morphology became same like as the EDTA capped ZnO nanoflower but the changes was in their distance between two neighbour flowers. Here the internal distance between two adjacent flowers were very less and the entire structure became more compact **(Fig 3H).** Here in this process chitosan biopolymer appeared as macromolecules which again aids in floral conformation of the ZnO nanomaterial over the N95 mask surface. Based on the SEM images, the ZnO nanomaterial were detected as flower like structures with zig-zag branches like pattern with flaky morphology with huge compactness [32]. The percentage of loading of variable compact ZnO nanoflower is determined by measuring the initial mass of the clean, untreated polypropylene fabric following with *in situ* synthesis and immobilization process of nanostructures and finally measure the mass of the Nano-treated, washed, and dried fabric. The loading percentage of Glycerol, PEG, EDTA and Chitosan assisted ZnO nanoflower tethered N95 mask is 38.7%, 54.8%, 77.4% and 87% respectively.

The room temperature fluorescence spectra of variable morphological ZnO nanoflowers prepared in different surface assisted medium measured at an excitation wavelength of 325 nm and the spectra were collected in the emission wavelength range of 350 nm to 600 nm using 1 cm path quartz cell were explained in **Supplementary 3 (S3).** The band gap energy of bulk ZnO material was about 3.4eV and due to this reason, the excitation energy used here was 3.8eV which was higher than the bulk one. Therefore, it was easy for an electron to be directly excited and jumped to the conduction band from the valence band. Additionally, the excitation of the deep level bands within the fixed band-gap was also possible. The fluorescence spectra of ZnO nanomaterials were exhibited two emission bands: one was in the UV region (392 nm) and the other was in the visible region (400nm - 650 nm). For glycerol assisted ZnO nanoflower sample the emission peak at 392 nm in UV region corresponded to the near bandedge (NBE) emission of ZnO and was due to the radiative recombination of free excitons. The violet emission peak at 427 nm was attributed to the formation of Zn interstitial defects and the green fluorescence at 536 nm may have originated from the antisite defects. It was well known that visible fluorescence was mainly due to defects related to deep-level emission, such as Zn interstitials and oxygen vacancies. PEG, EDTA and chitosan assisted all samples, the peak found at 392 nm and 427 nm (glycerol assisted sample) diminished and there a new band was created near 404 nm to 408 nm range which corresponded to the deep level violet emission band. In EDTA assisted sample another new band was found near blue-green emission zone at 465 nm wavelength which corresponds to the formation of oxygen vacancy defects. In all surface assisted nanomaterial the weak green fluorescence band observed at 536 nm can be related to recombination of electrons in the singly ionized oxygen vacancies with photo excited holes in the valence band. The intensity of green fluorescence band became lesser in PEG and glycerol assisted nanoflower whereas EDTA and chitosan produced more intensified green emission band. From the as mentioned fluorescence spectra results it was speculated that during the nanoflower formation the pattern or orientation of the floral petals produced changeable emission band. Except the glycerol assisted sample, all other surface modified samples showed five to six times more intensified blue emission band which causes formation of Zn interstitial defects. Upon changes the floral petals pattern with variable surface assisted molecules, the Zn interstitial defects became enhanced where the occupancy of the atoms in an interstitial site in the ZnO crystal structure became more which again confirms the presence of more impurities [33][34][35].

DLS spectra depicts the average particle size of differential ZnO nanoflower which was explained in **supplementary 4 (S4)**. From the DLS data it was indicted that ZnO nanoflowers were distributed in the range of 20-500 nm. The hydrodyanamic diameter of the ZnO nanoflowers was quite larger than the size of the nanomaterial reported from XRD, UV-Vis spectroscopy or SEM study was due to the fact that the size obtained from the DLS is always higher in value with respect of original material diameter because of the bounded ligand over the material surface and solvated water molecule.

### Antibacterial assay determination by both Broth Dilution assay and Kirby-Bauer test [36], [37], [38]

Minimum inhibitory concentration assay is a technique of the standard micro broth dilution assay recommended by the National Committee for Clinical Laboratory Standards (NCCLS), which has been developed to determine the antimicrobial activities of molecules where the visible growth of microorganisms in the agar broth have been noticed. There is an another assay termed as Kirby-Bauer test for antibiotic susceptibility (also called the disc diffusion test) is a standard test which is used to determine the resistance or sensitivity microbes to specific chemicals or molecules which can then be used by the clinician for treatment of patients with bacterial infections. The presence or absence of an inhibitory area termed as “inhibition zone” around the disc identifies the bacterial sensitivity to the drug. 100 µl (from log phase of bacterial culture) of both the *E.coli* (gram negative) and *B.subtilis* (gram positive) bacteria was spreaded over the agar plate and the uniform sizes of cups were digged out from it. Different ZnO nanoflower molecules along with two marketed antibiotic drugs termed as Amoxicillin and Erythromycin were poured into the cups with variable concentrations. If the organism is killed or inhibited by the concentration of the antibiotic or the nanomaterial there will be no growth in the immediate area around the cup which is called the zone of inhibition.

Various ZnO nanoflowers (PEG, glycerol, EDTA and chitosan assisted) in different concentrations i.e. 250 mg/ml, 1mg/ml, 750 µg/ml, 500 µg/ml, 250 µg/ml, 100 µg/ml and 1 µg/ml with adjusted bacterial concentration (10^8^ CFU/ml, 0.5 McFarland’s standard) for both gram negative *E.coli* and gram positive *B. subtilis* were used to determine MIC in 20 ml of LB broth. Simultaneously the marketed antibiotic drug named as Amoxicilin and Erythromycin was used as positive control, inoculated broth was treated as control and simple Deionised water was marked as negative control. These all culture tubes were incubated for 24 hr at 37 ° C in an orbital shaker at the speed of 200 rpm. The MIC value is the lowest concentration of ZnO nanomaterials where no visible growth is seen in the tubes. The visual turbidity of the tubes were noted, both before and after incubation to confirm the MIC value. Each 20 ml culture tube mixed with the different concentration compound was properly incubated in an orbital shaker at 37^0^C for 24 hrs and the corresponding optical density (OD) value as observed at 600 nm wavelength in a spectrophotometer. T_0_ value determined the initial OD value of the bacterial culture whereas T_24_ value determined the OD value of the bacterial culture after 24 hrs.

### Determination of MIC

Briefly overnight LB broth cultures of individual *E.coli* and *B.subtilis* bacteria (100 µl) were re-suspended in freshly prepared broth media (20ml) resulting in a suspension containing approximately 10^8^ CFU/ml. The initial OD of the bacterial culture media at 600 nm wavelength was observed by 0.01. To measure the MIC, 100 µl of bacterial solution was poured into the individual 20 ml culture broth. After that different concentration of 100 µl of individual ZnO nanoflower was added into the individual culture broth which was already occupied by 100 µl of bacterial culture solution. These all culture tubes were then incubated at 37°C for 24 h, and the ZnO-nanoflower concentration in the broth medium without visible growth of the bacterial cells was considered the MIC. A positive control contained LB Broth medium with tested bacterial concentrations and a negative control contained only inoculated broth were used in the study. The least concentration of individual ZnO nanoflowers that visually inhibit the bacterial growth after 24 hrs of incubation is defined as the MIC value of that particular molecules against that specific bacteria. Triplicate sets were performed to confirm the value for the tested bacteria. To determine the MBC, 100 µl of bacterial culture was taken from the treated molecules culture tubes that shows no turbidity/visibility under naked eye and streaked on LB agar plates. These all individual plates were kept in the incubator at 37^0^C for 24 hrs. The least concentration of the ZnO nanoflowers which prevented the growth of the bacteria can be determined as MBC.

In case of *E.coli* bacteria after 24 hr of incubation under aerobic conditions at 37°C, turbidity was noticed in the test tubes 1 µg/ml and 100 µg/ml containing individual ZnO nanoflower indicating the growth of bacteria. Whereas in concentrations of 250 µg/ml, 500 µg/ml, 750 µg/ml and 1 mg/ml, no turbidity was seen, exhibiting inhibition of bacterial growth. The suspension from these individual nanoflower treated tube was inoculated in LB agar plate and incubated for 24 hr to check the growth and no growth of bacteria was observed in all the concentrations hence confirming it as bactericidal. To study the MBC for different concentration of ZnO nanoflower when assessed for *E.coli* showed significant inhibition of growth for 250 µg/ml, 500 µg/ml, 750 µg/ml and 1 mg/ml, when compared to 1 µg/ml and 100 µg/ml and the MIC was obtained at 250 µg/ml. These results thus confirm that the MIC and MBC of ZnO nanoflowers for *E.coli* was found to be effective at dilution of 250 µg/ml. In addition the MIC and MBC of ZnO nanoflowers for *B. subtilis* was found to be effective at dilution of 100 µg/ml. **Fig 4A** revealed the sigmoidal growth curve of *E.coli* bacteria from initial hour to 24 hrs with one hr time interval. Additionally, how the bacterial growth was affected in presence of several ZnO nanoflowers, was also optimised in UV-Vis spectroscopy by studying the OD value at 600 nm wavelength and it revealed the fact that in presence of various nanoflower the bacterial growth was slow down or restricted. Among these four different ZnO nanoflowers the PEG assisted and glycerol mediated nanoflower with respect of other two i.e. chitosan and EDTA capped nanoflower showed more restricted bacterial growth. At T0, the OD value of E.coli shows 0.01 and at T24 it reaches the value of 1.08 which depicted the normal growth of the bacteria without any substance effect. The ZnO nanoflower concentration dependent OD values of E.coli was evaluated and it confirmed that near about 10-12 hrs the OD value of *E.coli* at 600 nm wavelength became very less and it reached the saturation point with compare to that of normal *E.coli* growth. But an interesting finding was observed for chitosan assisted ZnO nanoflower where the growth of the bacterial cell was sustained till 14 hrs and beyond that it reached the saturation point (**Fig 4B**). The growth of *B.subtilis* became more faster with quick cell doubling time with compare to that of *E.coli* (**Fig 4C**). Briefly overnight LB broth cultures of *B.subtilis* bacteria (100 µl) were re-suspended in freshly prepared LB broth media (20ml) resulting in a suspension containing approximately 10^8^ CFU/ml. The initial OD of the bacillus bacterial culture media was observed by 0.02 at T0 and after 24 hrs of incubation it reached the value of 2.98 which depicts the normal growth of the bacteria without any substance effect. In the case of *B.subtilis* bacteria, an interesting observation was found which was stated that for PEG and glycerol mediated ZnO nanoflower the growth of the bacteria was completely restricted at 8 hrs of incubation whereas this bacterial growth was till sustained upto 11 hrs of incubation for chitosan and EDTA assisted ZnO nanoflower **(Fig 4D**). After 24 hrs of incubation time the growth curve of both *E.coli* and *B.subtilis* bacteria along with the treatment of four different ZnO nanoflowers were depicted the fact that for PEG and EDTA assisted nanoflower sample the growth of *E.coli* bacteria became negligible with compare to that of other two nanoflower sample i.e. Glycerol and Chitosan mediated nanoflower.

Whereas in case of *B.subtilis* bacteria the PEG, EDTA and chitosan assisted nanoflower samples showed better growth inhibition property with respect of glycerol mediated nanoflower (shown in **Fig 5)**. The comparative analysis of both the *E.coli* and *B.subtilis* bacterial growth after 24 hrs in presence of differential concentration of ZnO nanomaterial and it postulated that the *B.subtilis* bacterial growth kinetic was much faster with compare to that of *E.coli* bacterial culture (without nanomaterial). While in presence of nanomaterial suspension, the growth kinetics of *B.subtilis* become less than the growth kinetics of *E.coli*. As we know the bacterial resistance was again used to determine the bacteriostatic activity of several ZnO nanoflowers against both *E.coli* and *B.subtilis* bacteria. LB agar was aseptically inoculated with both of the bacteria via the spread method. A hole which has the capacity of 100 µl liquid loading was created on top of the spread plate and there after loaded with the individual ZnO nanoflower sample with variable concentration. All the agar plates were kept in an incubator at 37^0^C for 24 hrs. After incubation all the culture plates were observed for “zone of inhibition” study which is a circular area relating to the antibacterial activity level upon the bacteria. Any zone of inhibition indicates the degree of sensitivity of the antimicrobial agent to that specific bacteria. The potency of the antimicrobial agent depends upon the zone periphery. On the contrary no “zone of inhibition” means total bacterial resistance towards the antimicrobial agent. Depending on the antimicrobial concentration the susceptibility of the bacteria can be determined by measuring the diameter of the inhibition zone from the centre of the hole and compare the value with standard one i.e the known marketed drug Amoxycilin and Erythromycin. The “zone of inhibition” study against *E.coli* bacteria by measuring the diameter around the hole and it was depicted in the **Fig 6A**. The “zone of inhibition” study resembles well with MIC study which stated that the PEG assisted nanoflower showed enhanced bacteriostatic activity upon decreased nanomaterial concentration and the highest inhibition zone diameter was seen at the concentration of 250µg/ml which was then gradually decreased. For glycerol assisted nanoflower sample the bacteriostatic behaviour was same as like as PEG assisted one. EDTA capped nanoflower, the bacteriostatic activity initially got declined upto the concentration of 500 µg/ml and there after it got suddenly elevated at the concentration of 250 µg/ml. **Fig 6B** depicted the “zone of inhibition” study against the bacteria *B.subtilis* by measuring the inhibition zone diameter and from the result it observed that the PEG and glycerol mediated nanoflower revealed enhanced bacteriostatic activity upon decreasing nanomaterial concentration and the concentration of 250 µg/ml showed greater inhibition zone diameter. Whereas the other two nanoflower revealed the viceversa i.e. the bacteriostatic activity tends to decrease upon decreasing nanomaterial concentration. As a result from bacteriostatic activity study it concluded the fact that not all nanoflower revealed the same zone of inhibition. The bacteriostatic activity in terms of the diameter of the inhibition zone also depended on the arrangement and orientation of the floral petals which thus helps to inhibit or restrict the bacterial growth.

**Fig 5:**
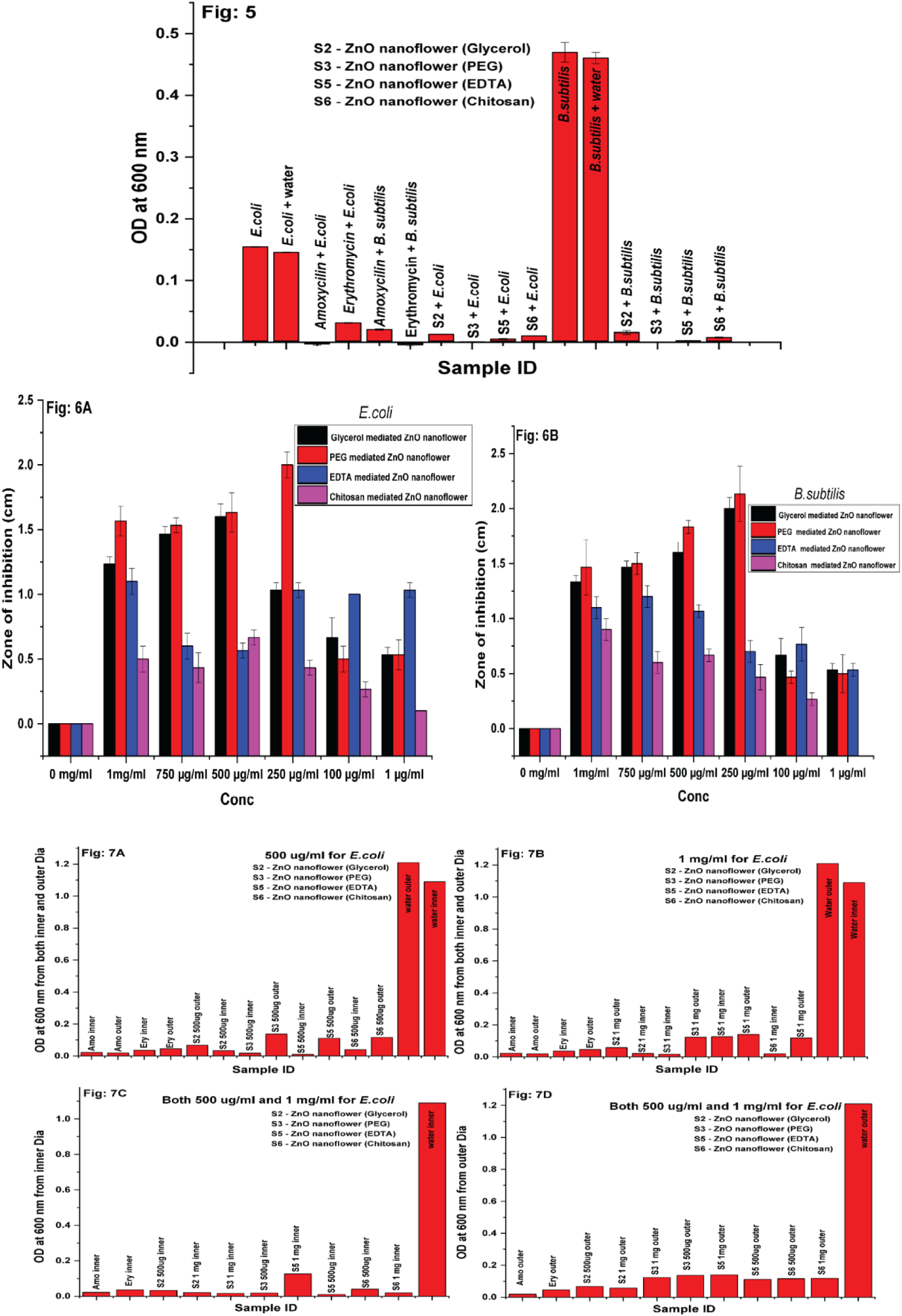
After 24 hrs of incubation time the growth curve of both *E.coli* and *B.subtilis* bacteria along with the treatment of four different ZnO nanoflower. **Fig 6A**: Zone of inhibition study against *E.coli* bacteria by using four different ZnO nanoflower (Kirbey Buyer assay); **6B:** Zone of inhibition study against *B.subtilis* bacteria by using four different ZnO nanoflower (Kirbey Buyer assay). Fig 7: Bactericidal activity study on a model bacteria (*E.coli)* by using modified Kirby Bauer assay.

To ensure the bactericidal activity of the bacteria, an interesting experiment was done. This experiment is the modified Kirby Bauer method where initially the bacteria was grown on the culture plate by spread method with created the hole to pour the drug molecules as well as nanomaterial samples later. After 24 hours of incubation period at 37°C in an aerobic condition the bacteria were completely grown over the agar plate. There after the hole which had the capacity of 100 µl loading was fully filled with individual ZnO nanoflower sample along with the two marketed drugs such as Amoxycilin and Erythromiocin. Here two concentration of material was taken i.e. the concentration where mostly the highest zone of inhibition was found from previously done Kirby Bauer test and the other concentration was the highest concentration. Again after 24 hrs of incubation period at 37°C the culture plate showed remarkable zone diameter around the hole. But here the difference is bacteria was already grown up over the culture plate and the zone diameter was found due to the molecule diffusion inside the agar. All the zone of inhibiton of Glycerol, PEG, EDTA and Chitosan assisted ZnO Nanostructures against both E. coli and B. subtilis bacteria have been depicted in **Supplementary figure 5 (S5)**. Now one loop full of bacteria was harvested from both the inner and outer side of the inhibition zone diameter and kept inside 10 ml of freshly prepared LB broth in a test tube to incubate for another 24 hrs at 37°C in an orbital shaker at 200 rpm. Finally, the OD values of all individual tubes were measured at 600 nm in an UV-Vis spectroscopy to compare the growth of the bacteria in different conditions. This test assay is the modified version of basic Kirby Bauer assay because in standard Kirby Bauer assay the concentration of the antimicrobial molecules can be determined for bacteriostatic activity against a specific bacteria where the bacteria can be grown over the culture plate along with the treatment of the antimicrobial components. For this reason, the antimicrobial components can inhibit the growth of the bacteria thus provides the information of bacteriostatic activity. Whereas this modified version of the Kirby Bauer test, initially the bacteria can grow full fledge over the culture plate and there after the antimicrobial components can be poured into the hole. Here in this case the antimicrobial components cannot inhibit the growth of the bacteria but it may bind to the bacterial cell and disrupt the outer membrane hence it completely kill the bacteria. **Fig 7(A-D)** depicted the fact that the variable surface modified ZnO nanoflowers can not only be treated as bacteriostatic, but it may also serve as bactericidal which was concluded by modified Kirby Bauer assay method which was done by the model system i.e. *E.coli* bacterial cell. Moreover among these all nanoflowers, EDTA and PEG assisted nanoflowers revealed best bactericidal effect against *E.coli* bacteria. Whereas EDTA, PEG and glycerol mediated nanoflowers showed best bacteriostatic activity against both *E.coli* and *B.subtilis* bacteria which was observed by standard Kirby Bauer test. To be precise, the antibacterial activity is governed by three major interconnected mechanisms: Reactive Oxygen Species (ROS) generation, Zinc ion (Zn^2+^) release, and direct mechanical damage. In general, the petal-like structure of the differential ZnO nanoflower structures exhibit high specific surface area, maximizing the contact points between the ZnO nanostructures and the biological/chemical environment (e.g., bacterial cells or water molecules). SEM study already revealed the floral structure formation is basically emerged from assembling of multiple nanorods. The interfaces and edges where the individual nanorods assemble are often rich in surface defects (like oxygen vacancies). These defects act as active sites for the generation of reactive species and the adsorption of reactants. The combination of high surface area and active defect sites gives ZnO nanofloral structure high antibacterial efficiency. ZnO is a wide-bandgap (∼3.37 eV) semiconductor. When ZnO absorbs a photon with energy greater than that of the band gap, an electron (e-) departs the valence band (VB) and jumps to conduction band (CB) with leaving behind a positively charged hole (h+) in the VB. The photogenerated e− and h+ migrate to the ZnO surface and react with adsorbed water (H_2_O) and molecular oxygen (O_2_), generating highly reactive oxygen species (ROS) thus initiates redox reaction. The generated ROS species are extremely destructive, causing oxidative stress and leading to lipid peroxidation by damaging the cell membrane, protein denaturation, and ultimately, DNA damage within the microbial cell, consequently it induces cell death. In the medium of the cell or moisture on the cell surface, the ZnO nanostructure undergoes slowly dissolution thus releasing toxic Zn^2+^ ions. Once inside the microbial cell, the Zn^2+^ ions interfere with essential biological processes such as enzyme inhibition and nutrient transport. Zn^2+^ ions have strong affection towards the SH-sulfhydryl groups, present in the respiratory chain of the bacterial enzyme, leading to their inactivation and the disruption of cellular respiration. On the other hand, the ions disrupt the cell’s proton motive force and overall transport system, inhibiting nutrient uptake thus leading to cell death due to nutrient deficiency. The sharp edges of ZnO nanofloral petal can cause abrasive damage or disruption to the cell membrane integrity, causing leakage in intracellular contents and rapid cell death. From the chemistry point of view, ZnO (especially at slightly acidic pH) can be positively charged, attracting and binding to the negatively charged microbial cell surface (due to components like teichoic acids in Gram-positive bacteria), increasing the local concentration of ZnO and Zn^2+^ [39].

### Conclusion

In summary, four different patterned ZnO nanoflowers with oriented attachment of floral petals were fabricated over the surface of N95 facial mask. It had been found that several surface modifier molecules such as PEG, EDTA, Glycerol and Chitosan and the thread of the mask was used as the growth enhancer of the ZnO nanomaterial in an oriented way. The synthesized ZnO nanoflowers were successfully characterized by various techniques which assured the nano-size and different patterned floral morphology of the ZnO. ZnO nanoflower materials were subjected as bacteriostatic as well as bactericidal against both *E.coli* and *B.subtilis*. Among the four different ZnO nanoflowers, PEG and EDTA assisted samples were found to be more antimicrobial property against both of these bacteria. The MIC and MBC value of ZnO nanoflowers for *E.coli* was found to be effective at dilution of 250 µg/ml whereas for *B. subtilis* it was found to be effective at the dilution of 100 µg/ml. Additionally it was concluded the fact that the variable surface modified ZnO nanoflowers can not only be treated as bacteriostatic, but it may also serve as bactericidal which was investigated by self-designed modified Kirby Bauer assay method. Ultimately The findings of the current study proposed that the ZnO nanoflower embedded N95 facial mask have a strong capacity for future applications in antimicrobial mask fabrication which is not only inhibit bacterial growth but it may also damage the bacterial cell partially.

## Acknowledgements

The authors were grateful to Mr. Gaurab Kumar Roy, a technical staff from Bose Institute for his help rendered in carrying out the SEM characterisation. We are also thankful to Mr. Dwaypayan Ghosh, JRF from Bose Institute for putting up his wonderful comments related to the upgradation of the work in the manuscript and constant moral support. Authors were also grateful to SERB, DBT India and Bose Institute for providing necessary support required for the above work.

## Author Contributions

Dr. Priyanka Bhadra is the first author of this manuscript who has conceptualised the work, invented the entire workflow design, writing – original draft, writing – review & editing, Validation, Supervision, Software, Project administration, Resources, Investigation, Formal analysis, Data curation and completely involved with the work protocol (both Nano and Bio part) with all of the characterisations thoroughly.

Rebanta Roy is the second author of this manuscript who has involved with entire antibacterial work, writing – original draft, writing – review & editing, investigation, Software, formal analysis and data curation. Also, since scratch point, he has attached with the first author with valuable discussions to strengthen the manuscript.

Prof Subhrangsu Chatterjee is the corresponding author who has scrutinized the entire work thoroughly, conceptualization, formal analysis and supervision of the entire work.

## Supplementary Figures

**S1:**
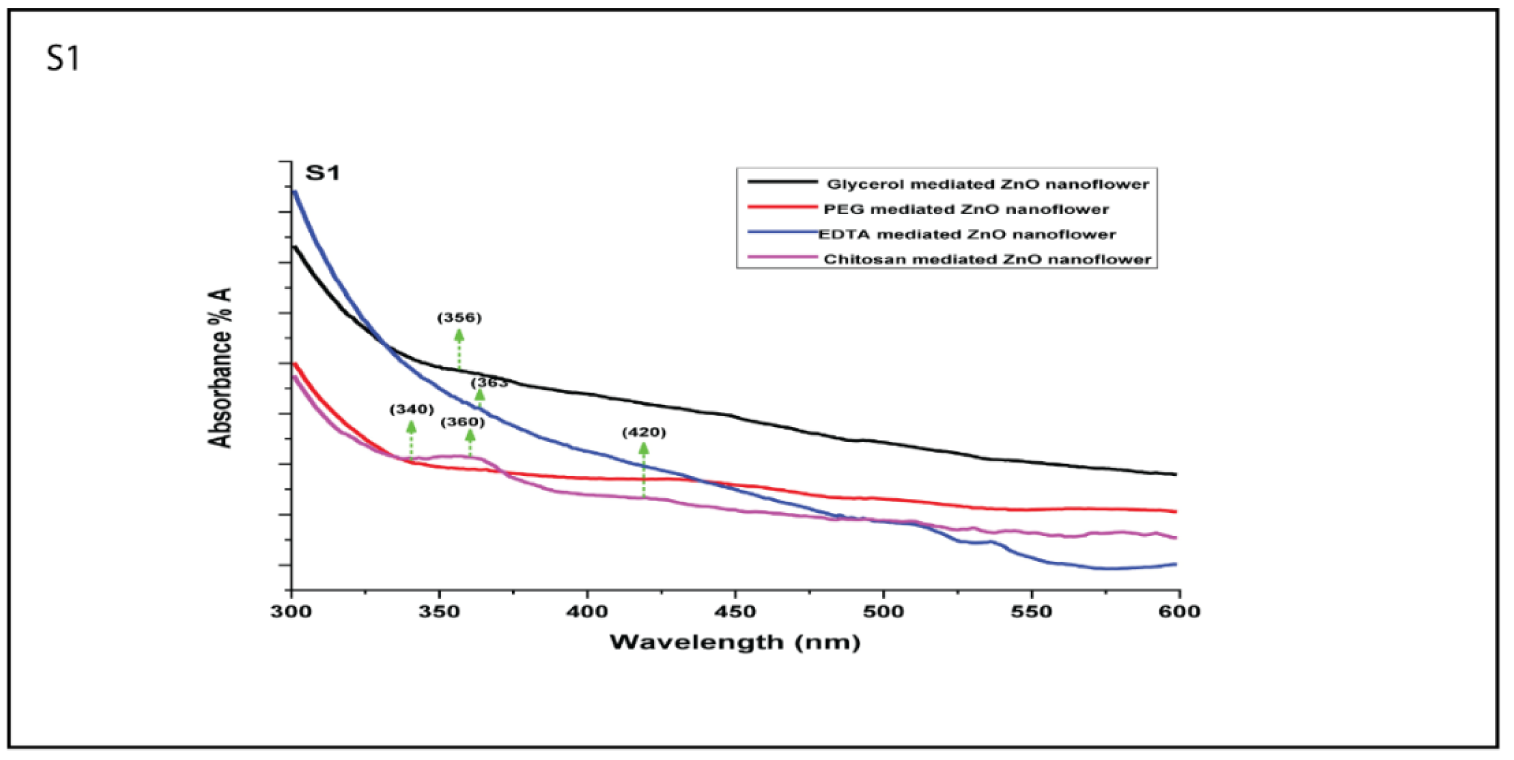
The UV-VIS absorption spectra of four different patterned ZnO nanoflower; **Sample 1 (black colour)** denotes the glycerol mediated ZnO nanomaterial shows continuous broad spectrum of absorption with a maxima at 356 nm. **Sample 2 (PEG assisted in Red colour)** and **Sample 3 (EDTA assisted in Blue colour)** shows the absorption maxima at 340 nm and 363 nm. Whereas the chitosan assisted **Sample 4 (Purple colour)** indicates a different kind of absorption spectra where the absorption maxima show at two variable positions such as 360 nm and 420 nm respectively.

**S2:**
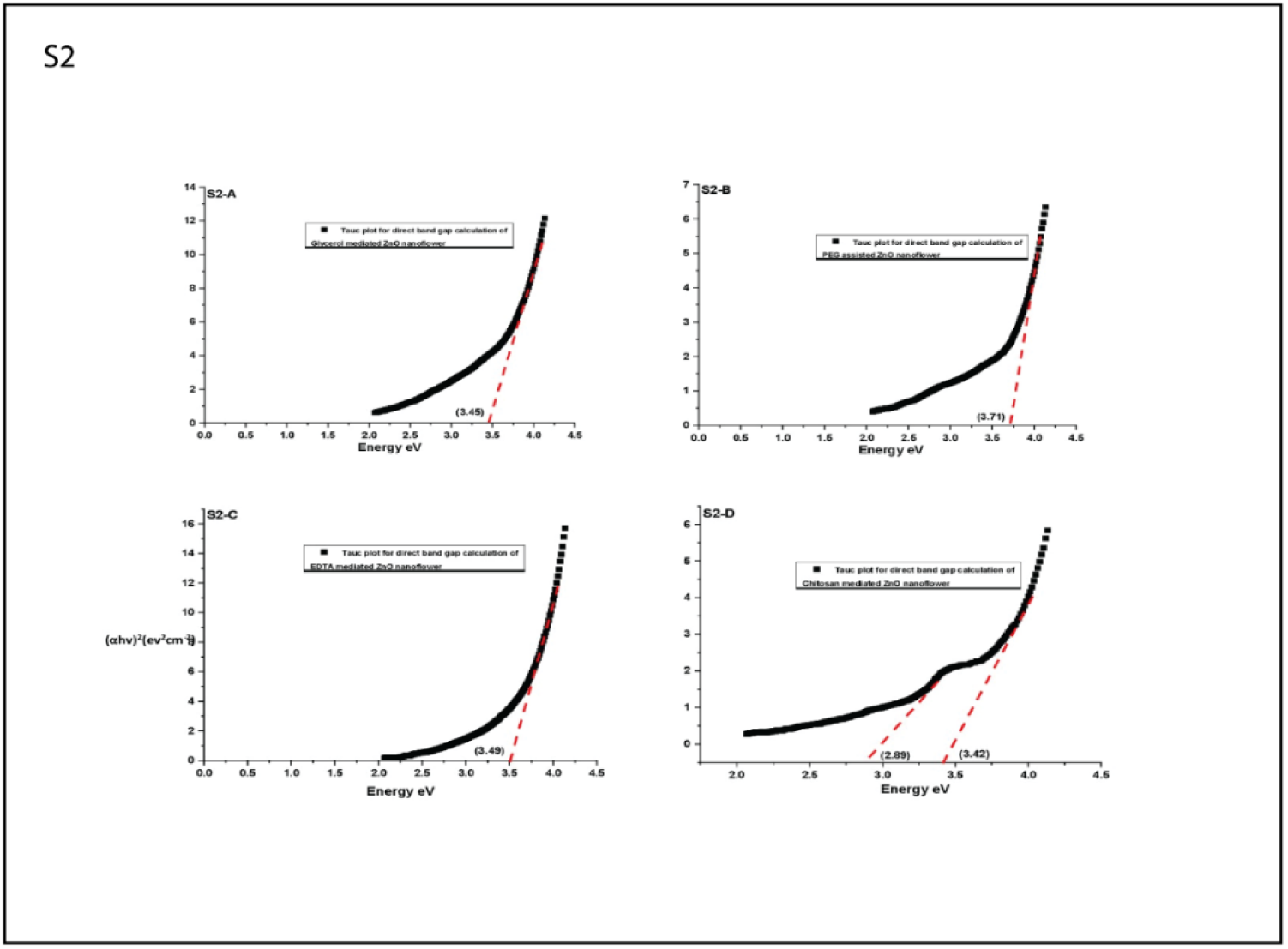
Band gap calculation from UV-Vis spectra by using Tauc plot; The band gap energy estimated from the absorption spectrum of four different ZnO nanomaterial was ∼3.45 eV **(S2-A for Glycerol assisted ZnO Nanoflower)**, ∼3.71 eV (S2-B for PEG assisted ZnO Nanoflower), ∼3.49 eV (S2-C for EDTA assisted ZnO Nanoflower) and ∼3.42 eV along with ∼ 2.89 eV (S2-D for Chitosan assisted ZnO Nanoflower)

**S3:**
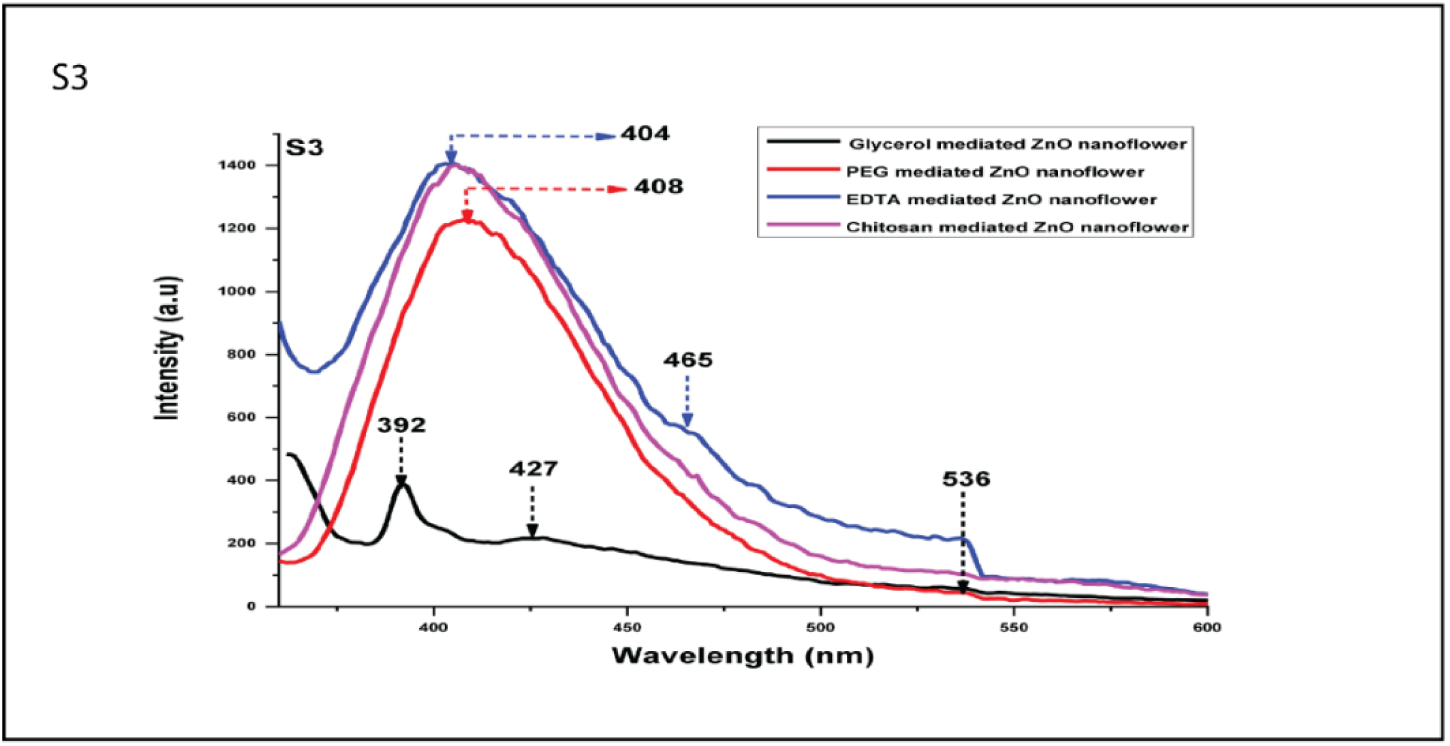
Room temperature fluorescence spectra of variable morphological ZnO nanoflowers prepared in different surface assisted medium; For **Glycerol assisted ZnO nanoflower** sample the emission peak observed at 392 nm in UV region and 427 nm in visible region denoted as violet emission peak; **PEG, EDTA and chitosan assisted all samples**, the peak found near 404 nm to 408 nm range and weak green fluorescence band observed at 536 nm.

**S4:**
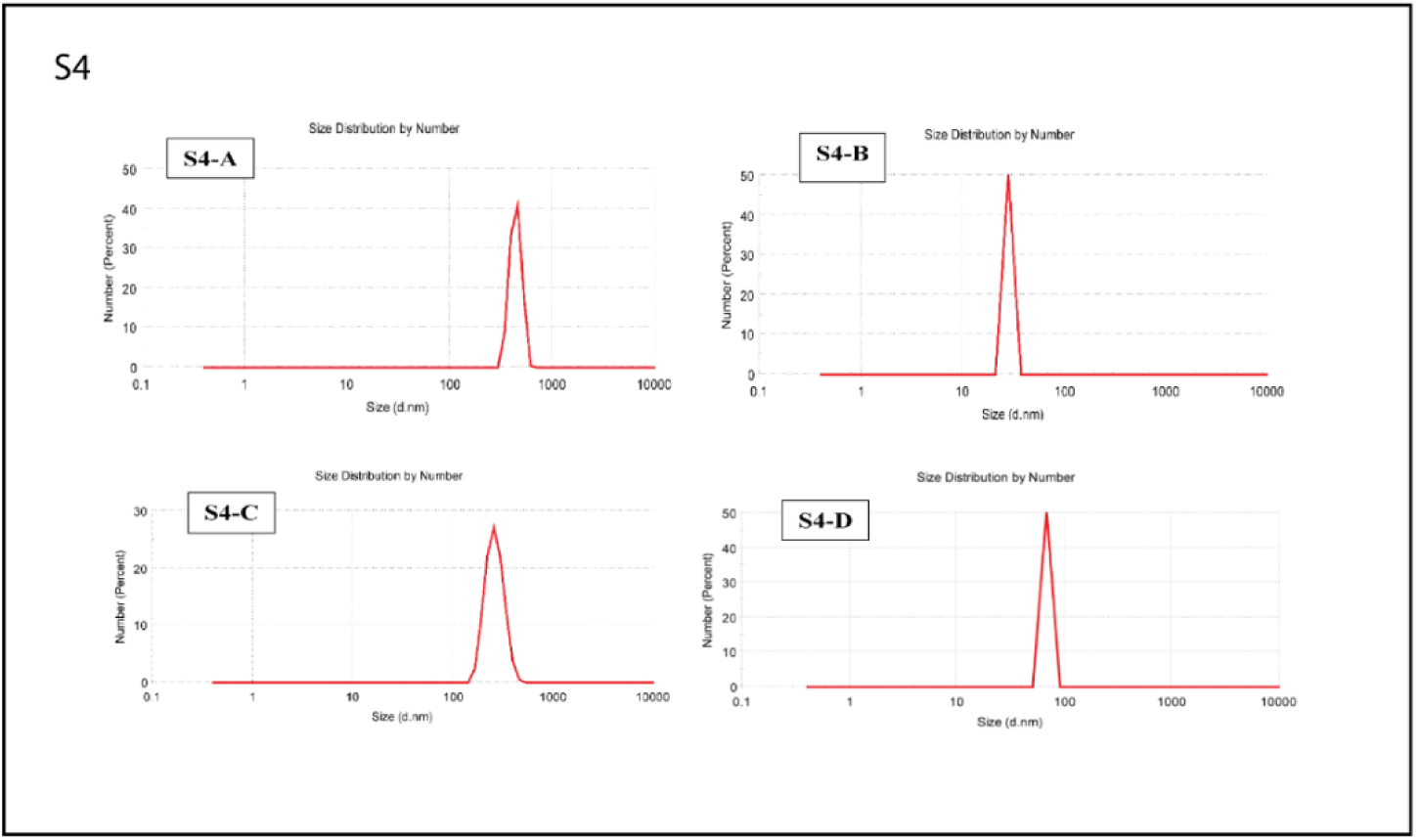
Average particle size of differential ZnO nanoflower determined by DLS. **A**. Glycerol mediated ZnO nanoflower, **B.** PEG mediated ZnO nanoflower, **C.** EDTA mediated ZnO nanoflower, **D**. Chitosan mediated ZnO nanoflower.

**S5(A-F):**
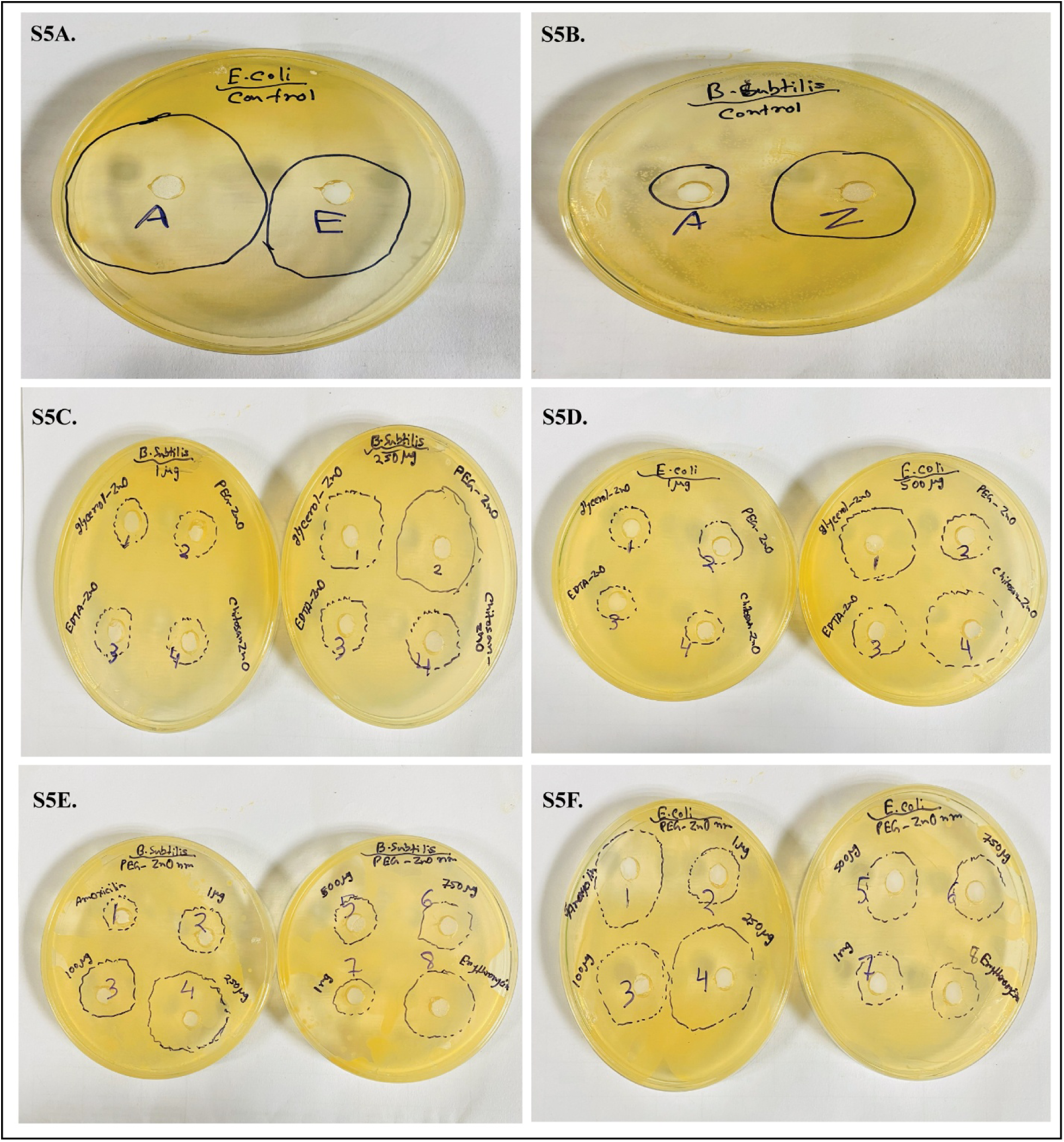
Zone of inhibition study of Glycerol, PEG, EDTA and Chitosan assisted ZnO Nanostructures against both *E. coli* and *B. subtilis* bacteria.

