## Supplementary Data for "De-Novo Designed Antibacterial N95 Facial Mask: Comprising a Nano-Garden Using ZnO Nanoflower"

**Supplementary Figure Caption**

**S1:** The UV-VIS absorption spectra of four different patterned ZnO nanoflower; **Sample 1 (black colour)** denotes the glycerol mediated ZnO nanomaterial shows continuous broad spectrum of absorption with a maxima at 356 nm. **Sample 2 (PEG assisted in Red colour)** and **Sample 3 (EDTA assisted in Blue colour)** shows the absorption maxima at 340 nm and 363 nm. Whereas the chitosan assisted **Sample 4 (Purple colour)** indicates a different kind of absorption spectra where the absorption maxima show at two variable positions such as 360 nm and 420 nm respectively

**S2:** Band gap calculation from UV-Vis spectra by using Tauc plot; The band gap energy estimated from the absorption spectrum of four different ZnO nanomaterial was ∼3.45 eV **(S2-A for Glycerol assisted ZnO Nanoflower)**, ∼3.71 eV **(S2-B for PEG assisted ZnO Nanoflower),** ∼3.49 eV **(S2-C for EDTA assisted ZnO Nanoflower)** and ∼3.42 eV along with ∼ 2.89 eV **(S2-D for Chitosan assisted ZnO Nanoflower)**

**S3:** Room temperature fluorescence spectra of variable morphological ZnO nanoflowers prepared in different surface assisted medium; For **Glycerol assisted ZnO nanoflower** sample the emission peak observed at 392 nm in UV region and 427 nm in visible region denoted as violet emission peak; **PEG, EDTA and chitosan assisted all samples**, the peak found near 404 nm to 408 nm range and weak green fluorescence band observed at 536 nm

**S4:** Average particle size of differential ZnO nanoflower determined by DLS. **A**. Glycerol mediated ZnO nanoflower, **B.** PEG mediated ZnO nanoflower, **C.** EDTA mediated ZnO nanoflower, **D**. Chitosan mediated ZnO nanoflower

**S5 (A-F):** Zone of inhibition study of Glycerol, PEG, EDTA and Chitosan assisted ZnO Nanostructures against both *E. coli* and *B. subtilis* bacteria.

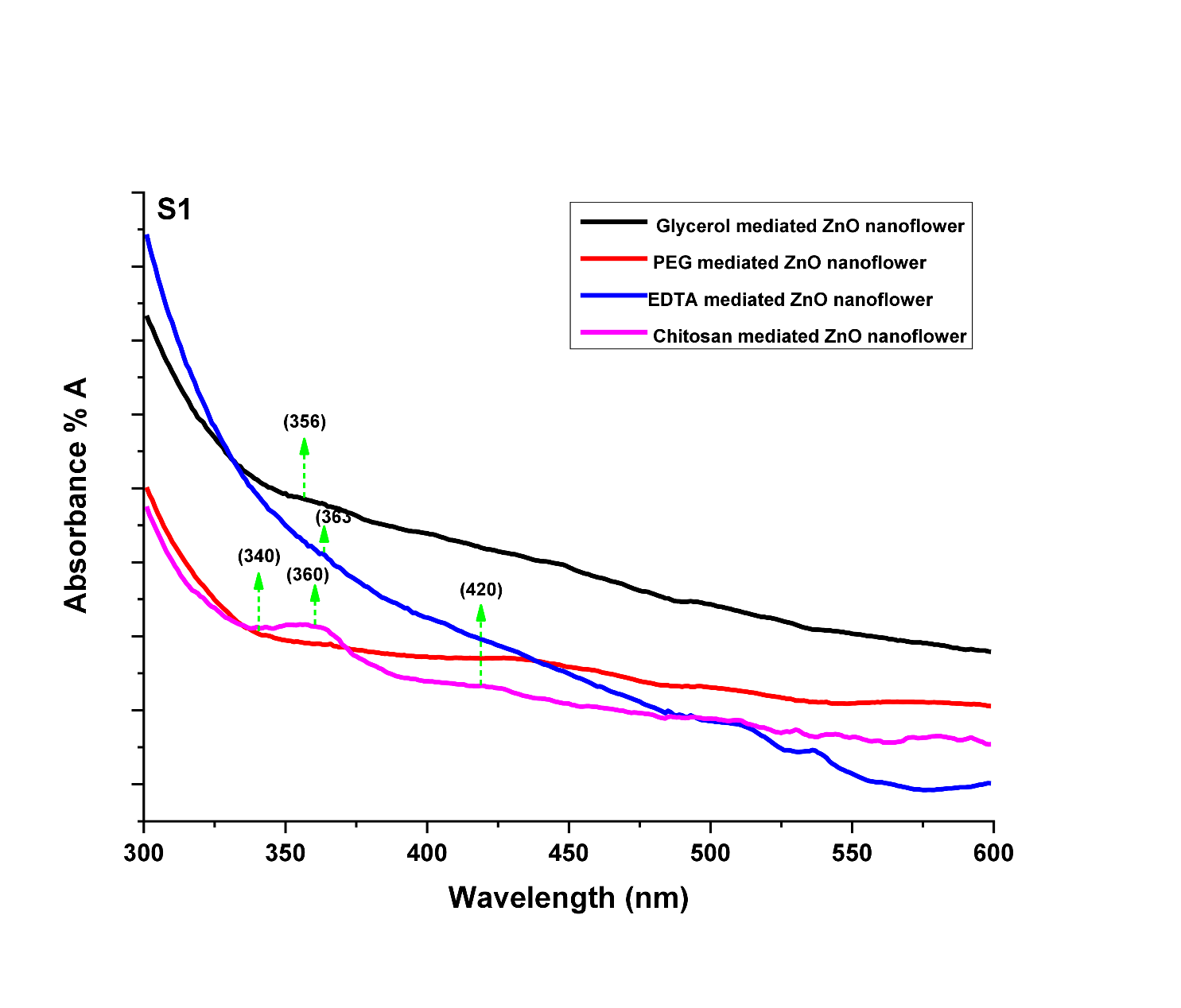

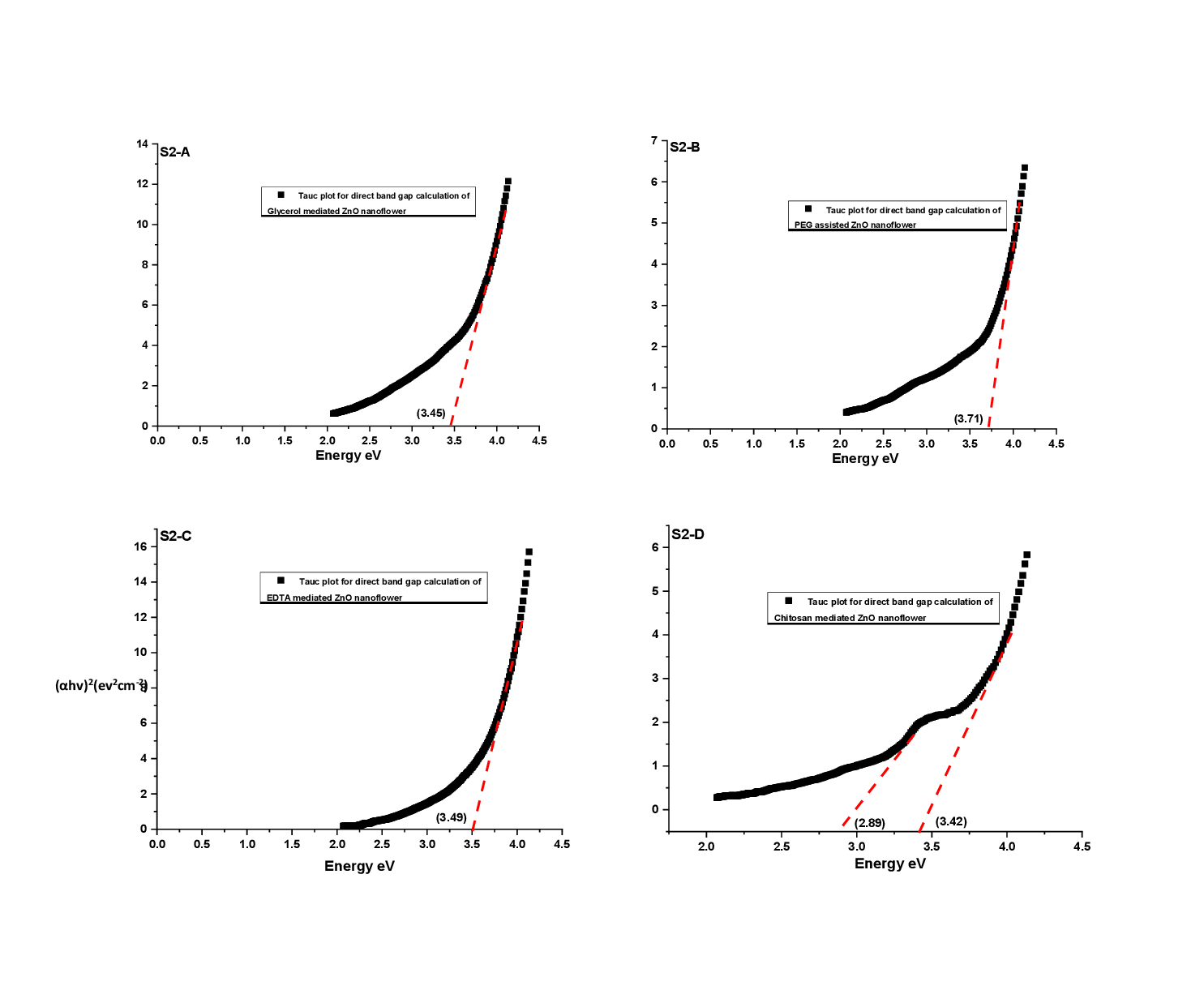

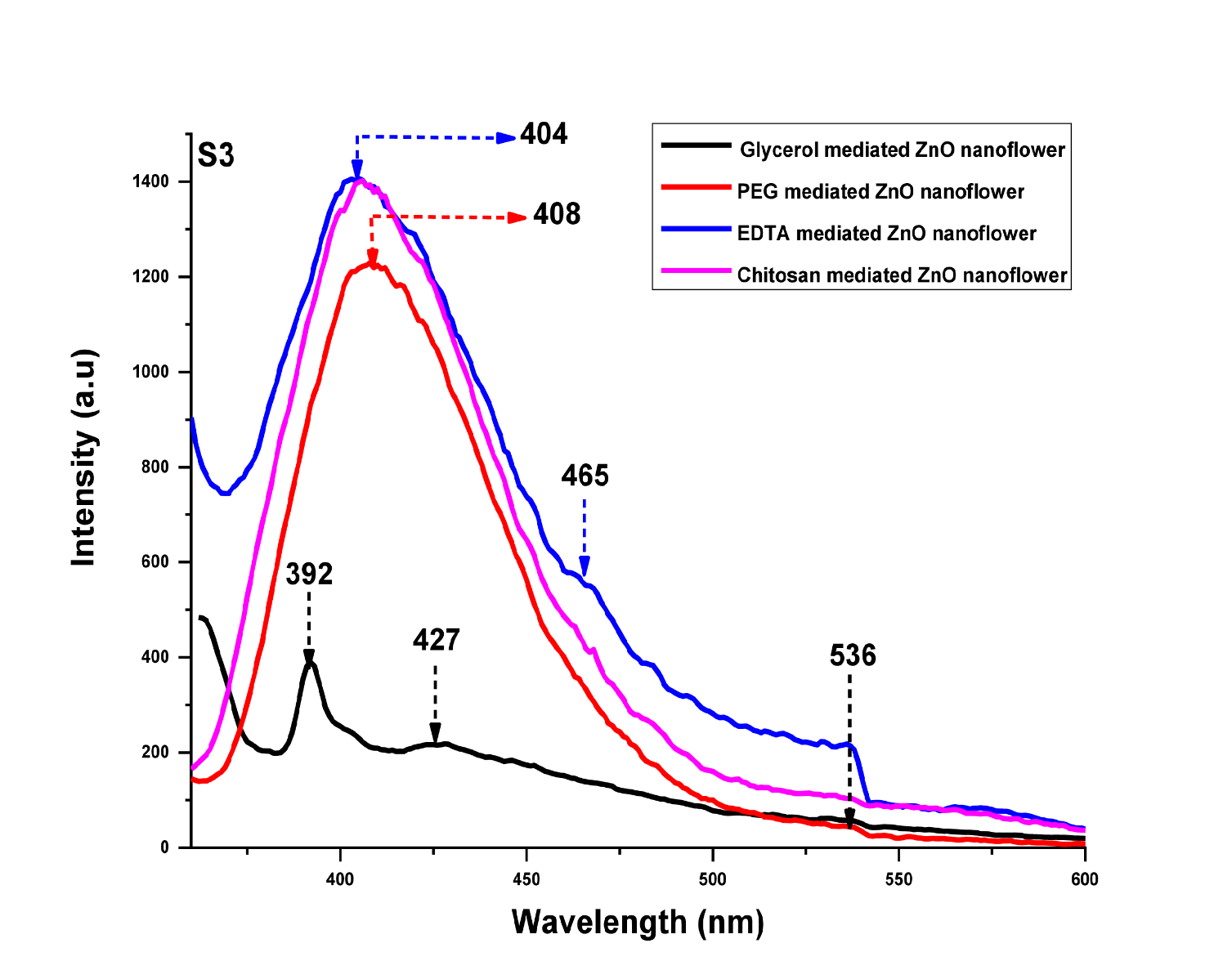

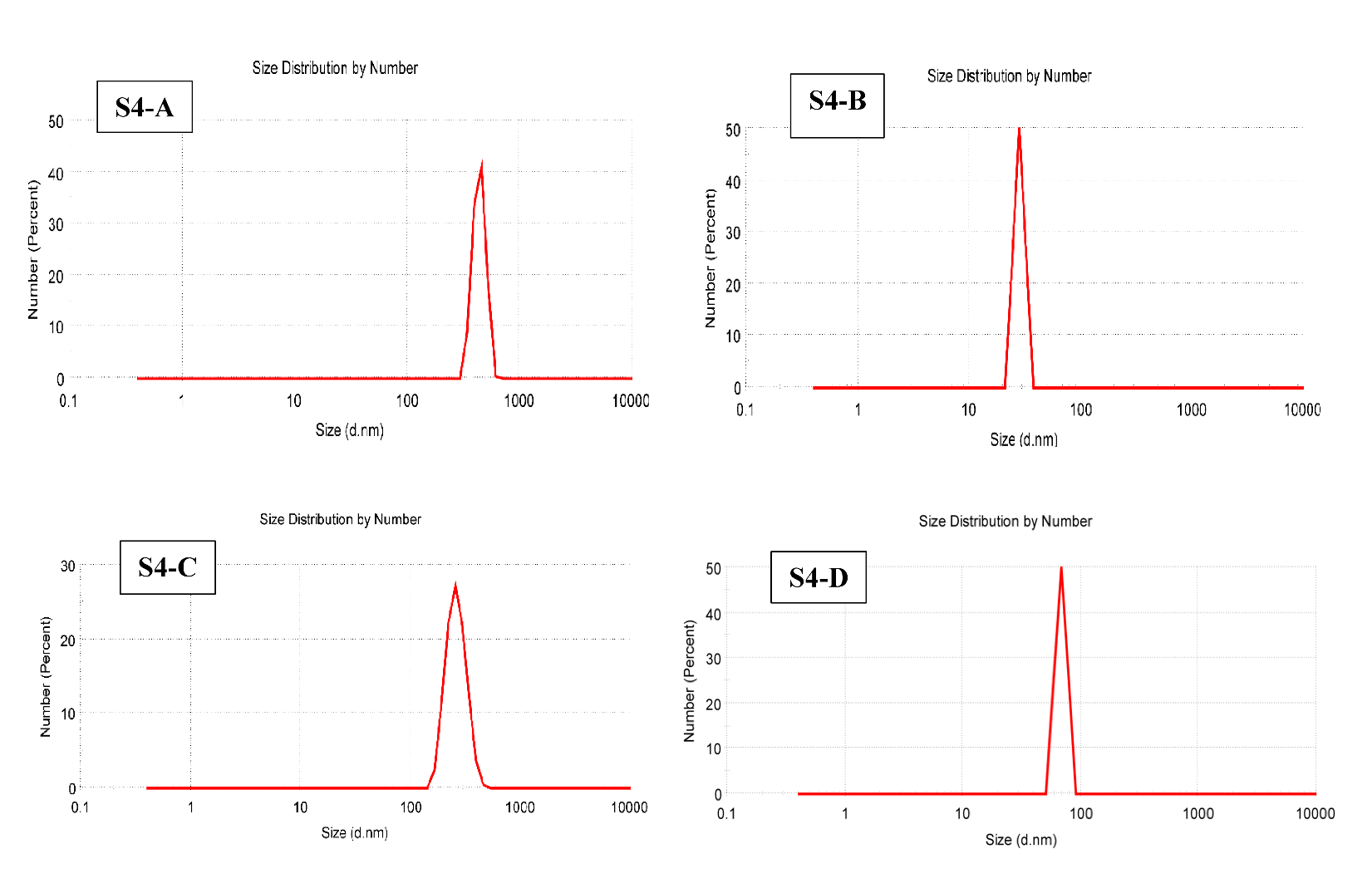

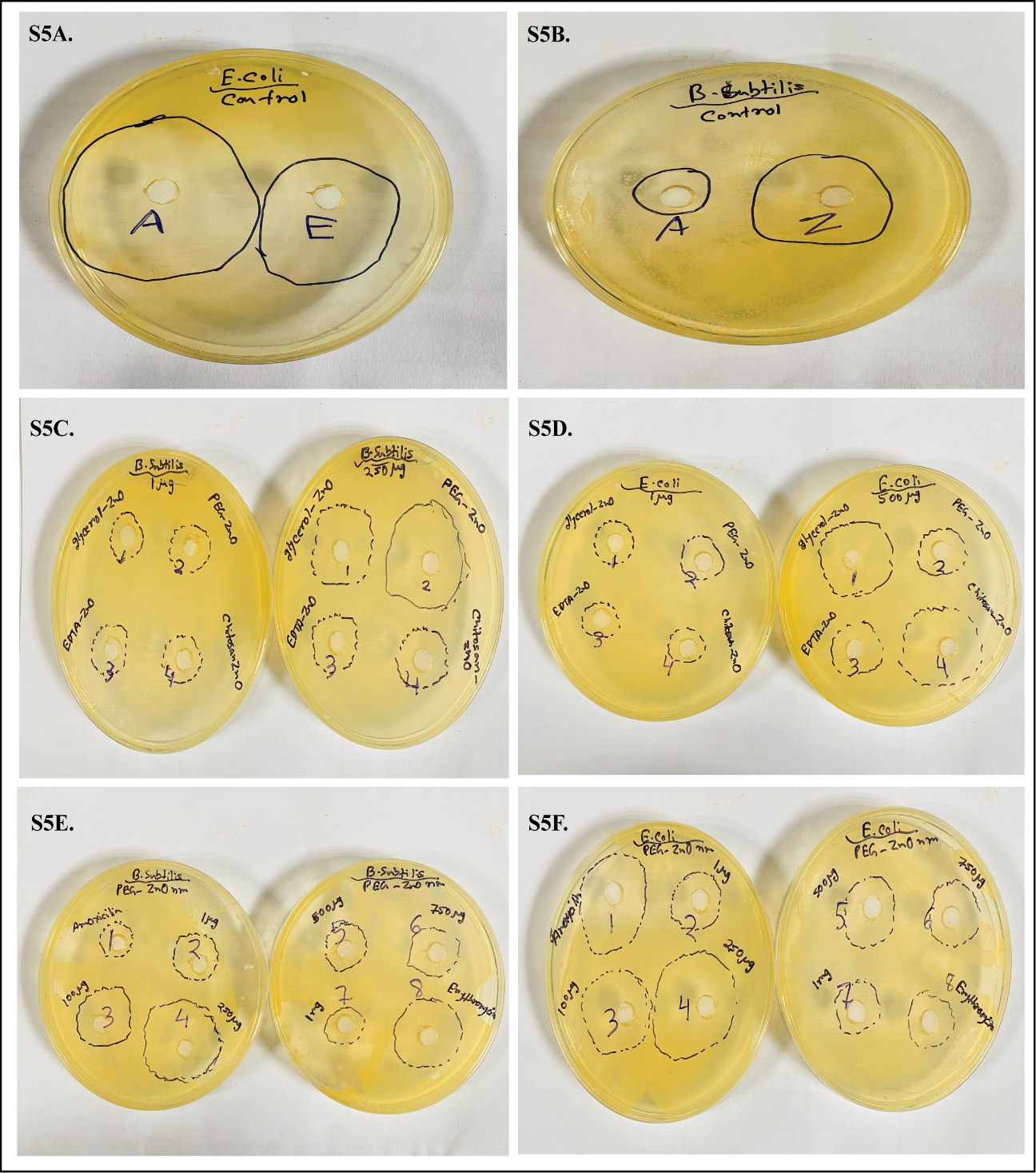
